# Programming bacteria for multiplexed DNA detection

**DOI:** 10.1101/2022.03.10.483875

**Authors:** Yu-Yu Cheng, Zhengyi Chen, Xinyun Cao, Tyler D. Ross, Tanya G. Falbel, Briana M. Burton, Ophelia S. Venturelli

## Abstract

DNA is a universal and programmable signal of living organisms. Here we developed cell-based DNA sensors by engineering the naturally competent bacterium *Bacillus subtilis* (*B. subtilis*) to detect specific DNA sequences in the environment. The DNA sensor strains can identify diverse bacterial species including major human pathogens with high specificity and sensitivity. Multiplexed detection of genomic DNA from different species in complex samples can be achieved by coupling the sensing mechanism to orthogonal fluorescent reporters. We also demonstrate that the DNA sensors can detect the presence of species in the complex samples without requiring DNA extraction. The modularity of the living cell-based DNA sensing mechanism and simple detection procedure could enable programmable DNA sensing for broad applications.

## INTRODUCTION

Next-generation engineered bacteria hold tremendous promise for a wide range of applications in human health, environment and agriculture by sensing key environmental signals, performing computation on these signals to regulate a response that modulates specific environmental parameters^1^. Developing specific and selective sensors of key environmental signals is a critical feature of next-generation engineered bacteria. For example, bacteria have been engineered to detect physical and chemical signals such as light, ultrasound, and quorum-sensing molecules^2–4^. These signals can be exploited to control the collective growth or gene expression of the bacterial population or mediate interactions between constituent community members^4–6^. In addition, we can exploit their sensing ability to achieve real-time monitoring of natural environments. For example, synthetic genetic circuits have been designed in *Escherichia coli* (*E. coli*) to sense signals produced by pathogens and use this information to regulate the production of antimicrobials that inhibit the target pathogen^7,8^. However, there are limited well-characterized and orthogonal signals that can be exploited to sense different bacterial species in a microbial community^9,10^.

DNA provides the blueprint for living organisms and is prevalent in natural environments^11^. Therefore, extracellular DNA (eDNA) could be exploited as a biomarker for identifying different species. Naturally competent bacteria have the ability to take up DNA from the environment and integrate imported sequences onto the genome based on sequence homology requirements. Horizontal gene transfer (HGT) via natural transformation has been shown to have a variety of benefits such as nutrient utilization, DNA repair, or acquisition of genes^12^. Since homologous recombination of imported DNA requires sequences of sufficient length and homology^13^, natural transformation could be exploited to build a selective cell-based DNA sensor.

We constructed a living cell-based DNA sensor by engineering the naturally competent bacterium *B. subtilis*. This circuit controls *B. subtilis* growth and fluorescence reporter genes in response to specific input DNA sequences. We demonstrate that the cell-based DNA sensor is sensitive and highly specific to species harboring the target DNA sequence. In addition, we demonstrate that our cell-based DNA sensor can perform multiplexed DNA detection in complex samples. The cell-based DNA sensors can detect DNA released from pre-treated donor cells (i.e. crude samples). Our detailed characterization of the cell-based DNA sensors *in vitro* provides a foundation for future *in vitro* DNA detection and *in situ* sense-and-respond DNA applications.

## RESULTS

### Construction of a living DNA sensor strain

To build the living cell-based DNA sensor, we exploited the natural competence ability of the well-characterized soil bacterium *B. subtilis*^14^. The natural competence ability of *B. subtilis* enables uptake of environmental DNA and integration of specific sequences with sufficient homology into genome via homologous recombination^15^. The efficiency of homologous recombination depends stringently on the sequence percent identity and length^13,16^, which can be exploited to build a highly specific DNA sensor.

To detect eDNA sequences in a programmable fashion, we constructed a synthetic genetic circuit in *B. subtilis* that implements a growth selection function based on the presence of target DNA sequence in the environment. The circuit consists of a xylose-inducible master regulator of competence *comK*^17^ and IPTG-inducible toxin-antitoxin system *txpA-ratA*^18^ and GFP regulated by the repressor LacI (**Fig. 1A,S1**). The target sequences were introduced to the flanking regions (upstream and downstream) of *txpA-ratA* and *lacI* and referred to as landing pads for homologous recombination.

**Figure 1.**
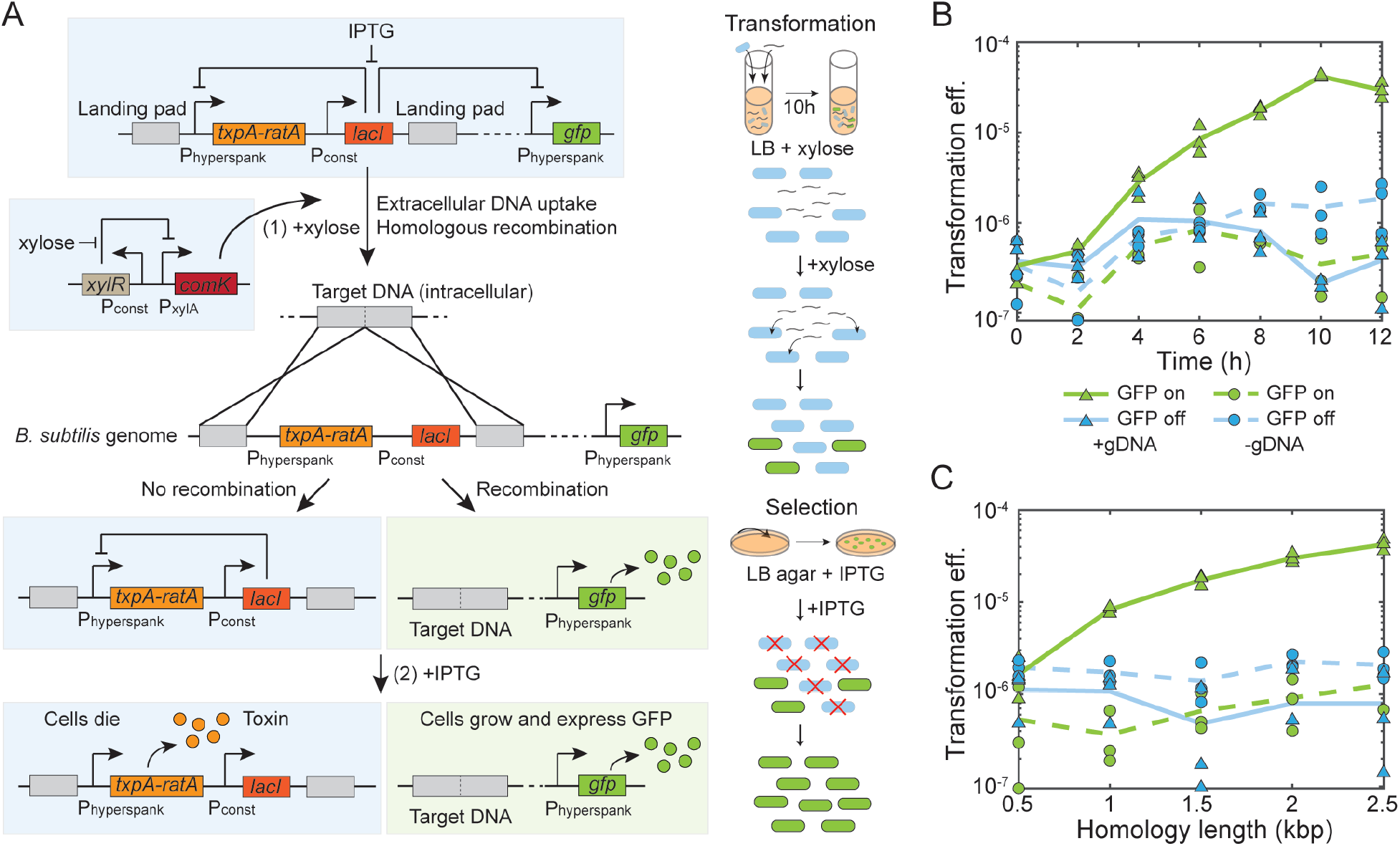
Construction and characterization of a living cell-based DNA sensor. **(A)** Schematic of a synthetic genetic circuit constructed in *B. subtilis* that allows the recognition of specific extracellular DNA. The xylose-inducible master regulator of competence *comK* can induce B. subtilis to take up eDNA and undergo homologous recombination. The toxin-antitoxin system *txpA-ratA* and a distal fluorescent reporter *gfp* are regulated by the repressor *lacI*. The target sequences are split into two and introduced to the flanking region of toxin and repressor as landing pad. In the presence of target DNA, homologous recombination can remove the toxin and the repressor, so a sub-population of transformed cells can express GFP. To select the transformed cells, IPTG can induce the toxin to kill the non-transformed cells, so the fluorescence can be enhanced by the growth of transformed cells. **(B)** Line plot of transformation efficiency versus transformation time for the *E. coli* DNA sensor in the presence (solid lines) or absence (dashed lines) of 100 ng/mL purified *E. coli* gDNA as input. The colonies on agar plate can be fluorescent (green) or not (blue), coming from background escape mutation or transformation. Transformation efficiency as output is optimal at 10 hr. Data points represent three biological replicates and lines are the average of the replicates. **(C)** Line plot of transformation efficiency versus homology length of *E. coli xdhABC* operon on each side of flanking region in the circuit in the presence (solid lines) or absence (dashed lines) of 100 ng/mL purified *E. coli* gDNA as input. The colonies on agar plate can be fluorescent (green) or not (blue). The transformation efficiency as output is above the background escape mutation when the homology length is equal or greater than 1 kbp and increases as the homology increases. Data points represent three biological replicates and lines are the average of replicates.

In the presence of xylose, ComK activates competence genes for DNA uptake and homologous recombination (**Fig. 1A**). Bistability and stochastic processes in the regulation of natural competence can yield a sub-population that can be transformed with extracellular DNA^19^. This naturally competent sub-population forms competence pili which bind to double stranded DNA outside the cell. The DNA is cleaved into single-stranded DNA (ssDNA) outside the cell membrane, and transported into the cell^12^. Inside the cell, RecA binds the ssDNA sequences and searches the *B. subtilis* genome for a region with sufficient homology. If the target DNA sequence is present, homologous recombination removes the toxin-antitoxin *txpA-ratA* and repressor *lacI*.In the presence of the chemical inducer Isopropyl β-D-1-thiogalactopyranoside (IPTG), cell growth and GFP expression is enabled in the transformed sub-population. Growth of the non-transformed subpopulation is inhibited by the activity of TxpA, which blocks cell wall synthesis^18^ (**Fig. 1A**).

We constructed a sensor for *E. coli* (EC sensor) by introducing the *xdhABC* operon onto the *B. subtilis* genome (landing pad region), which encodes genes for purine catabolism^20^. The *xdhABC* operon is a representative sequence that can detect a wide range of *E. coli* strains. This sequence is highly conserved such that 99% of 5000 *E. coli* genomes in the NCBI database contain this sequence with >95% coverage (the degree of alignment of the query sequence with a reference sequence) and >95% identity similarity (the percentage of bases that are identical to the target sequence within the aligned region).

To characterize the homology length needed for robust DNA sensing, we varied the homology length of the *xdhABC* operon in each landing pad (0.5 to 2.5 kb). We performed time-series measurements of transformation efficiency (number of colonies for transformed *B. subtilis* cells divided by the number total *B. subtilis* colonies) with 100 ng/mL *E. coli* genomic DNA (gDNA). The transformation efficiency is defined as the ratio of the number of transformed *B. subtilis* to the total number of *B. subtilis* based on colony forming units (CFU). Transformation efficiency plateaued at approximately 10 hr and the colonies expressed GFP (**Fig. 1B**). In addition, transformation efficiency increased with homology length at 10 hr (**Fig. 1C**). A homology length of 1 kb or greater was required to robustly sense the target sequence over the background frequency of escape mutants (10^-7^-10^-6^ frequency) that displayed heterogenous GFP expression (**Fig. S2A,B**). To achieve high performance of the DNA sensor (>10^2^ increase in transformation efficiency above background), we used a landing pad homology length of 2.5 kb (transformation efficiency of 10^-5^-10^-4^). In the transformed sub-population, homologous recombination was confirmed by sequencing to occur at the expected location with the elimination of *txpA-ratA* and *lacI* (**Fig. S2C,D**). The moderate number of escape mutants that displayed growth in the absence of gDNA had mutations in *txpA* or *lacI*, which reduced the growth inhibitory activity of TxpA (**Fig. S2E,F**). In sum, the synthetic genetic circuit enabled *B. subtilis* to sense specific DNA sequences present in the environment.

### Building living DNA sensors to sense human pathogens

Exploiting the modularity of the DNA sensing circuit, we replaced the landing pad region with specific sequences targeting different bacterial strains (**Fig. S1C**). To this end, we constructed DNA sensors to detect sequences harbored in human intestinal pathogens *Salmonella typhimurium*^21^ (*S. typhimurium*), *Clostridium difficile*^22^ (*C. difficile*), or the skin pathogen *Staphylococcus aureus*^23^ (*S. aureus*). We selected two 2.5 kb sequences in the pathogenicity island *sipBCDA* of *S. typhimurium* (ST sensor), the heme biosynthesis pathway *hemEH* in *S. aureus* (SA sensor), and the phenylalanyl-tRNA synthetase *pheST* in *C. difficile* (CD sensor)^24–26^.

The selected set of target DNA sequences are highly conserved within a given species (ST sensor: 94%, SA sensor: 96%, and CD sensor: 96% all with >95% coverage and >95% identity similarity). In addition, some of the sequences are linked to virulence activities of the pathogen or encode enzymes that are critical for fitness^24–26^. To further explore the conservation of the target sequences across different strains, we performed nucleotide BLAST using the NCBI Database to quantify the homology coverage and sequence similarity across species. The pathogenicity island *sipBCDA* in *S. typhimurium* was found only in *Salmonella enterica* species and infrequently observed in other species (**Fig. S3A**). Homologs in other species have low coverage and identity similarity, suggesting that the pathogenicity island could be a good target sequence for this species (**Fig. S3A**). The heme biosynthesis pathway *hemEH* in *S. aureus* and phenylalanyl-tRNA synthetase *pheST* in *C. difficile* are conserved in some closely related strains with varying degrees of similarity and coverage (**Fig. 2I,K**). The *E. coli* MG1655 *xdhABC* purine catabolism operon is found in other closely related bacteria such as *Shigella* with high coverage and identity similarity (**Fig. S3B**). Although the target sequences for building the different DNA sensor strains varied in the degree of specificity based on bioinformatic analyses, a detailed characterization of circuit performance could guide the design of optimized cell-based DNA sensors for future applications.

**Figure 2.**
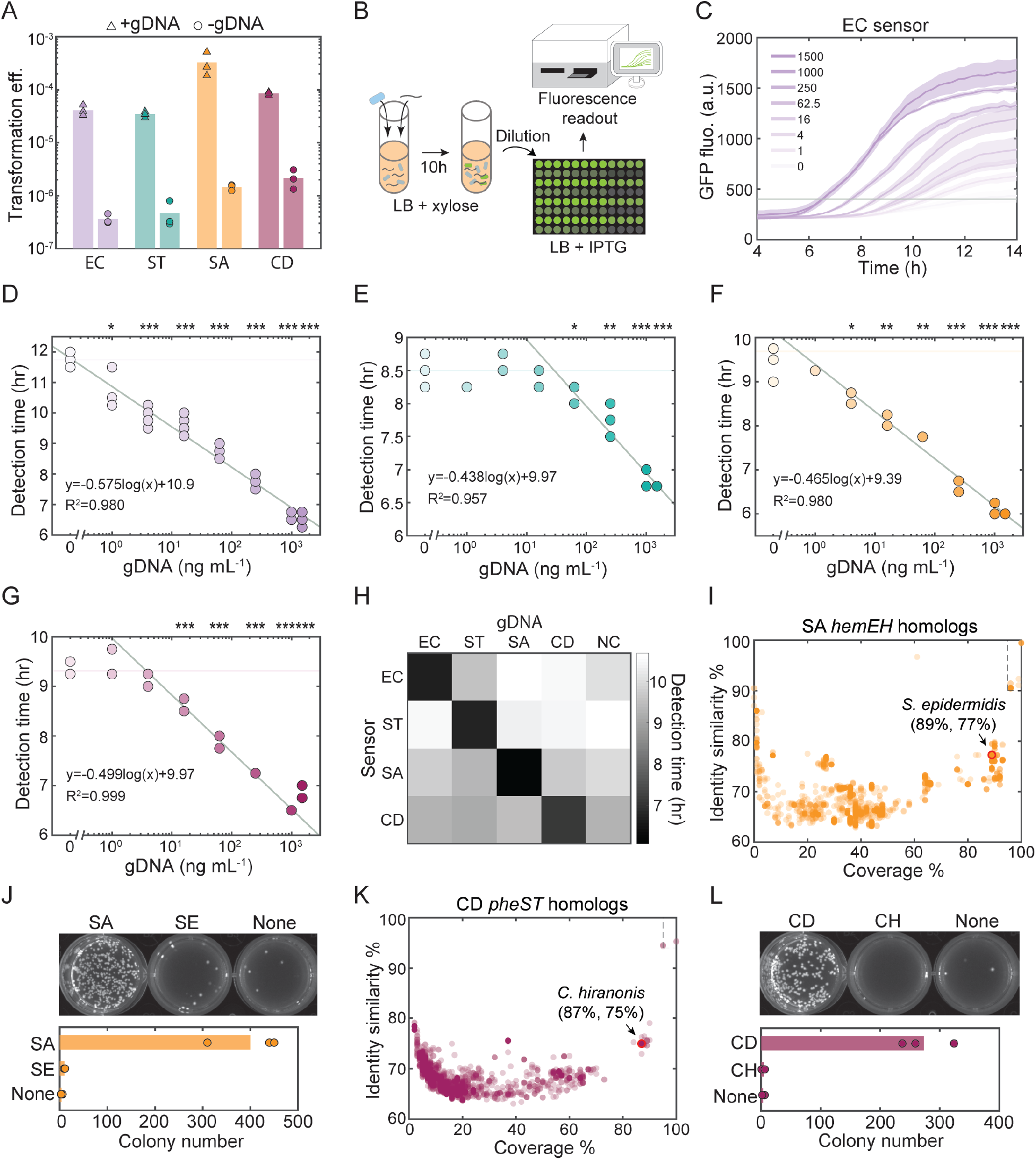
Cell-based sensors can detect DNA from diverse bacteria with high sensitivity and specificity. **(A)** Transformation efficiency of DNA sensors that can detect *E. coli* (EC sensor), *S. typhimurium* (ST sensor), *S. aureus* (SA sensor), or *C. difficile* (CD sensor) in the presence of 100 ng/mL extracted gDNA (triangles) or no gDNA (circles) at 10 hr. Bar represents the average of three biological replicates. **(B)** Schematic of experimental procedure for quantifying GFP expression after transformation. **(C)** Time-series measurements of GFP expression of EC sensor in liquid medium after the transformation of varying *E. coli* gDNA concentrations (ng/mL). A threshold of GFP 400 was used to determine the detection time for each gDNA concentration. Line is the average of four technical replicates and the shaded region represents one standard deviation from the average. Detection time versus gDNA concentration for **(D)** EC sensor, **(E)** ST sensor, **(F)** SA sensor, and **(G)** CD sensor. Horizontal line (pale color) is the background GFP fluorescence in the absence of gDNA. Unpaired *t*-test was performed to determine if the detection time with specific DNA concentration is different from the background fluorescence, and *, **, and *** denote *p*-values < 0.05, 0.01 and 0.001, respectively. A straight line was fitted to the transformation efficiencies versus logarithmic gDNA concentrations with statistical differences. The slope of the fitted line was determined by the cell growth of *B. subtilis*, while the intercept was determined by the background escape mutation. The coefficient of determination R^2^ shows the goodness of the fit. **(H)** Detection time of each DNA sensor after the transformation of 100 ng/mL gDNA of a given donor species or no gDNA representing the negative control (NC). Each sensor expressed GFP 3~4 hours earlier in the presence of gDNA from its target strain than gDNA from other strains. Data are the average of four technical replicates. **(I)** Nucleotide BLAST search of 5000 bp *S. aureus hemEH* in the NCBI database. Each circle represents a homolog found in species other than *S. aureus* and its coverage and identity similarity. A closely related human commensal strain *S. epidermidis* with high coverage and similarity was selected for specificity test using SA sensor. The region within dashed lines indicates *Staphylococcus* species that could be recognized by the SA sensor based on their high similarity of homologous sequences. **(J)** Comparison of colony numbers of SA sensor (with GFP expression) on selective agar plate after the transformation of 100 ng/mL *S. aureus* gDNA, 100 ng/mL *S. epidermidis* gDNA or no gDNA. Bar represents the average of three technical replicates. SA sensor can distinguish between *S. aureus* and closely related *S. epidermidis*. **(K)** Nucleotide BLAST search of 5000 bp *C. difficile pheST* in the NCBI database. Each circle represents a homolog found in species other than *C. difficile* and its coverage and identity similarity. A closely related human commensal strain *C. hiranonis* with high coverage and similarity was selected for specificity test using CD sensor. The region within dashed lines indicates *Clostridium* species that could be recognized by the CD sensor based on their high similarity of homologous sequences. **(L)** Comparison of colony numbers of CD sensor (with GFP expression) on selective agar plate after the transformation of 100 ng/mL *C. difficile* gDNA, 100 ng/mL *C. hiranonis* gDNA or no gDNA. Bar represents the average of three technical replicates. CD sensor can distinguish between *C. difficile* and closely related *C. hiranonis*.

The four sensors robustly detected the presence of 100 ng/mL target gDNA over background (0 ng/mL gDNA) based on transformation efficiency (**Fig. 2A**). We evaluated the sensitivity of each DNA sensor strain by performing time-series GFP measurements in liquid culture after being transformed with a wide range of gDNA concentrations (0-1500 ng/mL gDNA) from single species (**Fig. 2B,C,S4**). We evaluated the time required for each culture to display a fluorescence level higher than a threshold (i.e. detection time) (**Fig. 2C,S4**). The sensitivity of the circuit was evaluated as the lowest gDNA concentration that yielded a statistically significant difference in the detection time in the presence versus absence of gDNA (**Fig. 2D-G**).

The relationship between the log transformed gDNA concentration and detection time is linear due to the exponential growth of the fluorescent *B. subtilis* sub-population successfully transformed with the input target sequence (**Fig. S5**). Therefore, to assess the range of gDNA concentrations that can be accurately sensed, we inferred the parameters of a linear function fit to the log transformed gDNA concentration versus detection time (**Fig. 2D-G**). The inferred slope of the linear function is determined by the cell doubling time (~0.5 hour) and intercept is determined by the background mutation frequency (**Fig. 2D-G**). The EC, SA, and CD DNA sensor strains displayed high sensitivity of 1-16 ng/mL (10^5^-10^6^ chromosome copy number/mL), whereas the ST sensor displayed a lower sensitivity (62.5 ng/mL, 10^7^ chromosome copy number/mL). While the DNA sensor strains displayed lower sensitivities than quantitative real-time polymerase chain reaction (qPCR) reported for *E. coli* (3.5×10^3^ CFU/mL in pure culture^27^), the observed sensitivities are within the range of the sensitivities reported for the lateral flow immunoassay (1.8×10^5^ CFU/mL for a pure culture of *E. coli*^28^).

To characterize the specificity of each DNA sensor strain to the target sequence, we performed time-series fluorescence measurements in liquid culture in response to all individual species gDNA. The fluorescence signal was observed at a substantially earlier time (6.1-7.1 hr) in the presence of the corresponding species’ gDNA than in the presence of a non-target species’ gDNA or in the absence of DNA (9.1-10.7 hr) (**Fig. 2H, S6**). This demonstrates that the cell-based DNA sensors were highly specific to the target sequence.

In the context of microbial communities, the cell-based DNA sensors may need to distinguish between closely related species. Therefore, we evaluated the ability of the DNA sensors to distinguish between closely related species with similar target sequences. To this end, we measured the transformation frequency of the SA sensor in the presence of gDNA (100 ng/mL) derived from *S. epidermidis. S. epidermidis* is a closely related human skin commensal bacterium that harbors a similar *hemEH* sequence to the SA sensor landing pad region (89% coverage and 77% identity similarity) (**Fig. 2I**). Since the number of total *B. subtilis* colonies was similar in the presence and absence of gDNA, we quantified the number of transformed colonies as opposed to transformation efficiency (**Fig. 2A,S7**). The number of colonies in the presence of *S. epidermidis* gDNA (SE) was substantially lower than in the presence of *S. aureus* gDNA (SA) and similar to the absence of DNA (**Fig. 2J**). Similarly, we characterized the ability of the CD sensor to detect gDNA derived from a human gut commensal bacterium *C. hiranonis*, a close relative of *C. difficile* that contains a similar *pheST* sequence in its genome to the landing pad region in CD sensor (87% coverage and 75% identity similarity). The number of colonies in the presence of 100 ng/mL *C. hiranonis* gDNA (CH) was substantially lower than in the presence of *C. difficile* gDNA (CD) and also similar to the absence of DNA (**Fig. 2K,L**). These data demonstrate that the cell-based DNA sensors are highly specific to species that harbor an exact match to the target sequence. Therefore, the sensors do not display false positives in the presence of closely related species that harbor similar target sequences. The high specificity of the sensors is due to the stringent requirements for homologous recombination in *B. subtilis*^13,29^.

### Multiplexed detection of pathogen DNA in complex samples

Since certain future applications may require sensing of more than one organism, we tested the ability of the DNA sensors to detect more than one species within mixed DNA samples. To this end, we constructed individual sensors with orthogonal fluorescent reporters to achieve multiplexed DNA detection. Exploiting the modularity of the circuit, we constructed an RFP-labeled ST sensor (ST-R) and a BFP-labeled SA sensor (SA-B), in addition to the GFP-labeled EC sensor (EC-G) (**Fig. S1C**). We introduced gDNA (200 ng/mL) extracted from each of the three target strains into a culture containing EC-G, ST-R, and SA-B sensors and determined the number of fluorescent colonies for each reporter (**Fig. 3A**). The sensors accurately reported the presence/absence of all combinations of species’ gDNA reliably (**Fig. 3B,S8**). Therefore, a mixture of DNA sensor strains each individually labeled with a unique fluorescent reporter enabled multiplexed detection of gDNA derived from different species.

**Figure 3.**
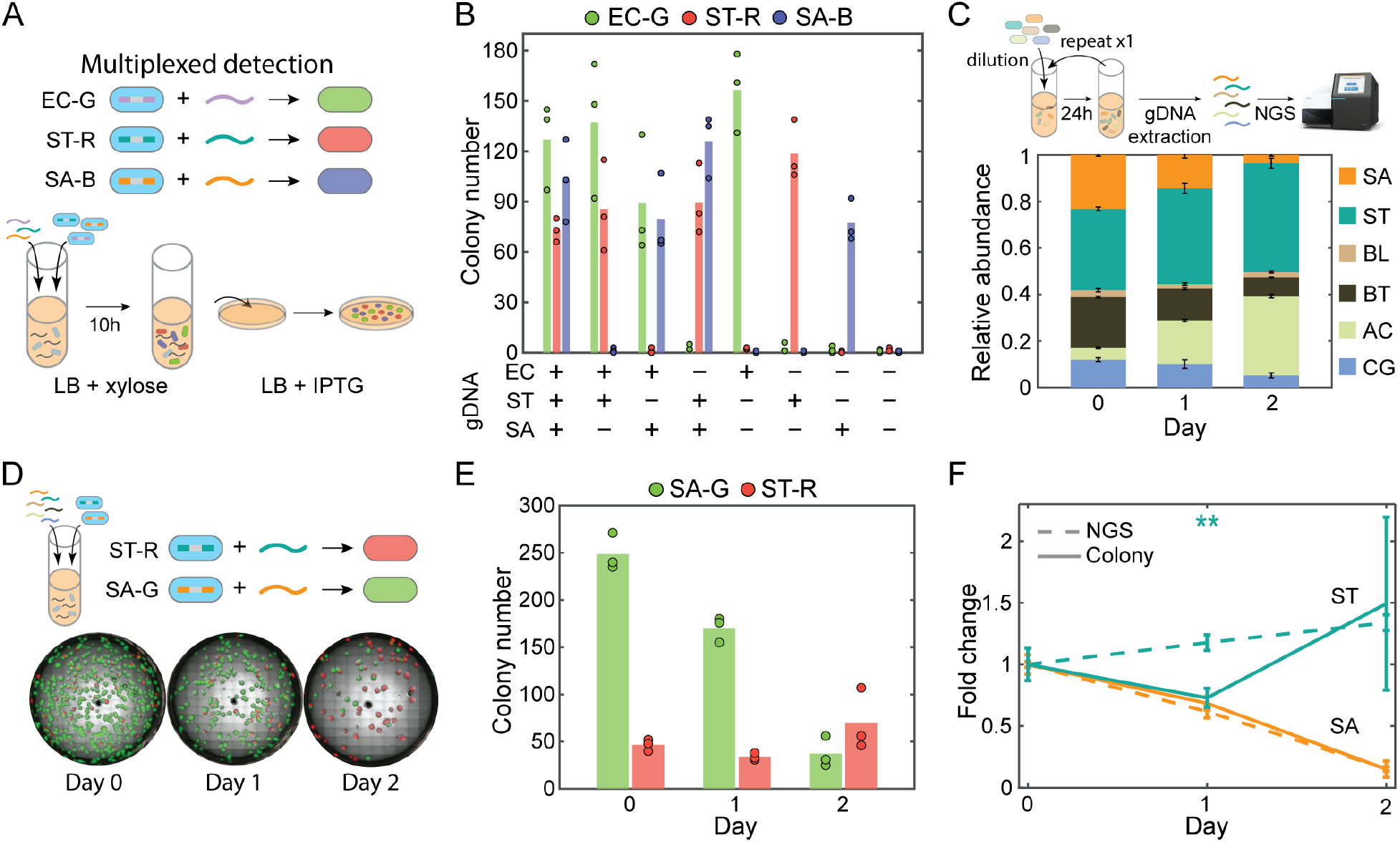
DNA sensors can perform multiplexed detection in complex DNA samples. **(A)** Schematic of experimental procedure using the living cell-based DNA sensors for multiplexed detection. *E. coli, S. typhimurium*, and *S. aureus* DNA sensors were labeled GFP (EC-G), RFP (ST-R) and BFP (SA-B) for detection, respectively. Three sensors and gDNA extracted from different strains were mixed into liquid medium for transformation. Transformed cells were selected on agar and colonies expressing GFP, RFP and BFP can indicate the presence of target DNA. **(B)** Numbers of GFP, RFP, or BFP-expressing colonies on agar plate after the transformation of different combinations of gDNA extracted from *E. coli*, *S. typhimurium*, or *S. aureus*. Sensors can report the presence of target DNA for all 8 different combinations. Bar represents the average of three technical replicates. **(C)** Relative abundance of six species in a synthetic gut microbial community composed of *S. aureus* (SA), *S. typhimurium* (ST), *Bifidobacterium longum* (BL), *Bacteroides thetaiotaomicron* (BT), *Anaerostipes caccae* (AC), and *Clostridium asparagiforme* (CG) over two days of growth. The six bacteria were co-cultured in liquid medium anaerobically for 24 hours and cell culture was diluted in fresh medium once for species to continue their competition. 16S rRNA gene of each strain was PCR amplified and sequenced using NGS to determine the relative abundance over time. Bar represents the average of three technical replicates of 16S rRNA sequencing. *S. typhimurium* abundance remained similar while *S. aureus* abundance decreased over time. **(D)** Representative fluorescence images of transformed SA-G and ST-R sensors on selective agar plate after the transformation of gDNA extracted from the bacterial community in a mixture of SA-G and ST-R sensors. The numbers of green and red colonies indicate the abundance of *S. aureus* and *S. typhimurium* in gut microbial community, respectively. **(E)** Numbers of GFP or RFP-expressing colonies on agar plate after the transformation of gDNA extracted from the community at different time. Green colonies (SA-G) decreased over time while red colonies (ST-R) remained similar numbers. Bar represents the average of three technical replicates. **(F)** Fold change of *S. aureus* and *S. typhimurium* abundance compared to day 1 characterized by NGS (dashed line) or cell-based detection method (solid line). Measurements of *S. aureus* (orange) by the two methods were similar, while characterization of *S. typhimurium* (green) by the cell-based detection method had larger variability than NGS and the data at Day 1 were statistically different determined by Unpaired *t*-test (*p*-values = 0.0015).

To investigate if multiplexed detection can be achieved for samples derived from a complex microbial community, we constructed a four-member human gut community composed of diverse commensal bacteria from three major phyla in human gut – *Anaerostipes caccae* (AC, Firmicutes), *Bacteroides thetaiotaomicron* (BT, Bacteroidetes), *Bifidobacterium longum* (BL, Actinobacteria), and *Clostridium asparagiforme* (CG, Firmicutes). This community also contained the target pathogens *S. typhimurium* (ST) and *S. aureus* (SA). We tested whether the DNA sensors could accurately report the relative abundance of the two pathogens during community assembly. The 6-member community was inoculated in equal initial species proportions based on absorbance at 600 nm (OD600, Day 0) and cultured anaerobically for 24 hr (Day 1). An aliquot of the community was transferred to fresh media and community composition was characterized following an additional 24 hr (Day 2). Based on 16S rRNA gene sequencing, the abundance of *S. typhimurium* was similar as a function of time whereas the abundance of *S. aureus* decreased over time (**Fig. 3C**).

We characterized the ability of the DNA sensors to accurately track the temporal trends in species abundance by introducing purified community gDNA collected at different times into a mixed culture of the ST-R and SA-G sensors (**Fig. 3D**). Due to the low abundance of target species in the sample, a higher amount of DNA (1 μg/mL) was used for transformation. Consistent with the trends based on 16S rRNA gene sequencing, the number of GFP fluorescent colonies of the SA-G sensor decreased at sequential time points, whereas the number of RFP fluorescent colonies of ST-R sensor were similar at sequential time points (**Fig. 3D-F**). The SA sensor displayed better performance in mirroring the trend from 16S rRNA gene sequencing than the ST sensor, consistent with its higher sensitivity than other sensors (**Fig. 2A,3E,F**). For the community lacking *S. typhimurium* and *S. aureus*, a much smaller number of background colonies was detected than in the 6-member community. This implies that the ST and SA sensors were specific to the target species gDNA and did not generate false positives in the presence of the other constituent community member gDNA (**Fig. S9**). In sum, our results show that accurate multiplexed DNA detection can be achieved in samples derived from multi-species microbial communities.

### Detection of target species without DNA extraction

Specific bacterial species have been shown to release eDNA in response to environmental stimuli^11^, suggesting that the DNA sensor could detect species without requiring prior gDNA purification. To test this possibility, we co-cultured individual DNA sensor strains with the corresponding donor species with an initial OD600 0.1 of the target strain (1.22×10^8^ CFU/mL, 1.07×10^8^ CFU/mL, 3.2×10^8^ CFU/mL, and 1.1×10^7^ CFU/mL for *E. coli, S. typhimurium, S. aureus, and C. difficile*, respectively) (**Fig. 4A**). Since the other species could compete with *B. subtilis*, we introduced specific antibiotics (ABX) to inhibit the growth of the donor cells and enhance donor eDNA release. The DNA sensor strains are resistant to the antibiotics since they harbor the appropriate antibiotic resistance genes. In the presence of 100 μg/mL spectinomycin, the DNA sensors displayed robust detection of *E. coli*, *S. typhimurium*, and *S. aureus* (**Fig. 4B**). The addition of spectinomycin was not required for *C. difficile* detection since the growth of *C. difficile* is negatively impacted by the presence of oxygen^30^.

**Figure 4.**
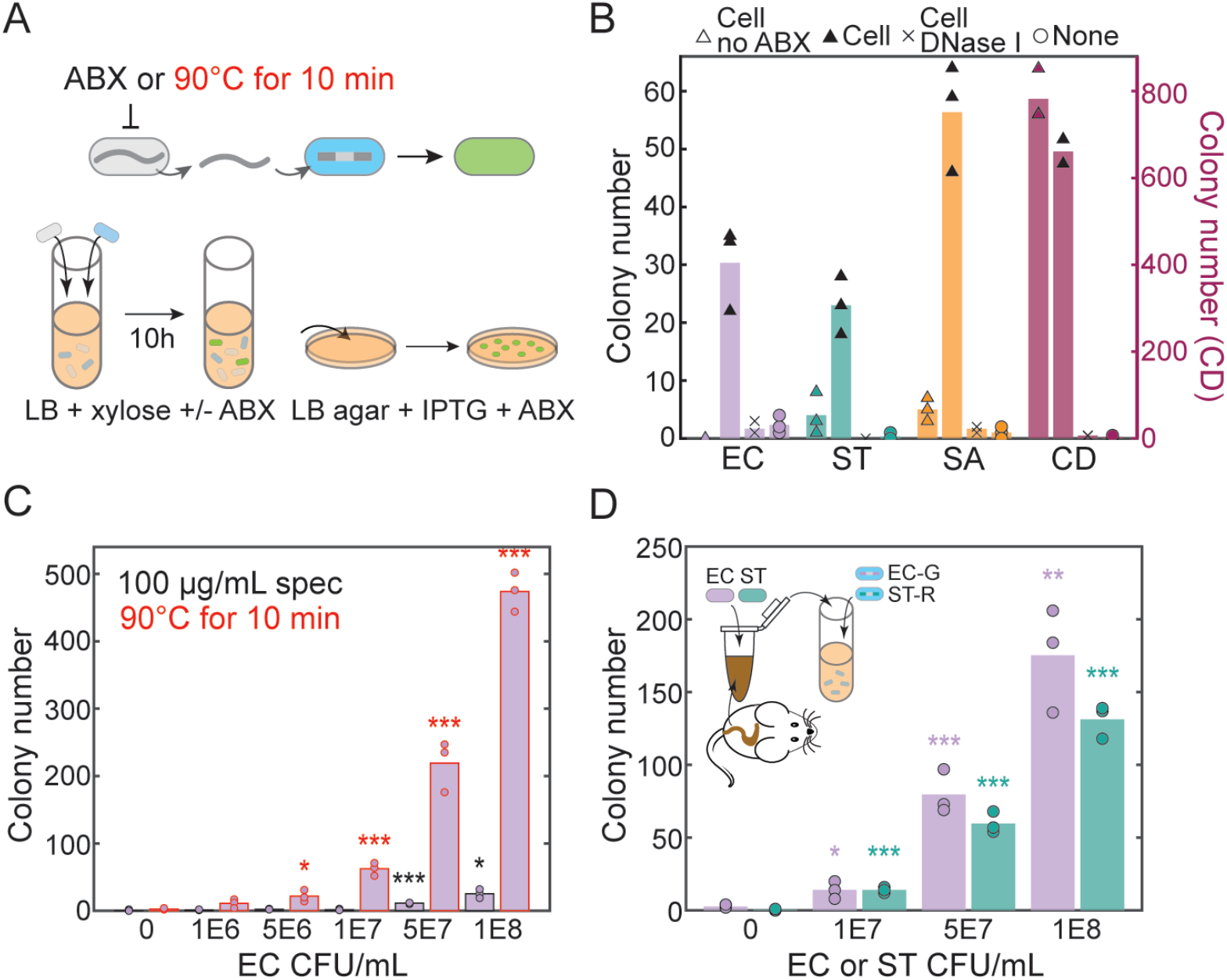
DNA sensors can directly detect target species without DNA extraction. **(A)** Schematic of experimental design for the detection in the co-culture. Sensor and target strain were co-cultured in liquid medium with or without selective antibiotics (ABX) to determine the effect of antibiotic for detection. Heat-killed target strain was also tested without the use of antibiotics in the co-culture. The cell culture was plated on IPTG and ABX (1 μg/mL erythromycin and 25 μg/mL lincomycin) agar plates to select for transformed *B. subtilis*. **(B)** Colony numbers of transformed EC, ST, SA, and CD sensors co-cultured with (1) target cell, (2) target cell and 100 μg/mL spectinomycin (spec), (3) target cell, spec, and 1 unit/mL DNase I, and (4) spec only. Spectinomycin can enhance the detection of *E. coli, S. typhimurium*, and *S. aureus*, but it is not required for *C. difficile* detection. Addition of DNase I in the co-culture reduced the numbers of transformed sensors significantly. **(C)** Colony number of transformed EC sensor co-cultured with *E. coli* using spectinomycin or co-cultured with heat-treated *E. coli*. Heat treatment of *E. coli* can significantly increase the number of transformed EC sensor. **(D)** Colony number of transformed EC-G and ST-R sensors co-cultured with heat-treated mice cecal samples spiked in with different amounts of *E. coli* and *S. typhimurium*. Both sensors can detect the presence of target strains in cecal samples. Unpaired *t*-test was performed to determine if the colony number is different from no cell condition, and *, **, and *** denote *p*-values < 0.05, 0.01, and 0.001, respectively. Bar represents the average of three technical replicates.

To confirm the transformation was mediated by the eDNA released from the target strain, DNase I (1 unit/mL) was added into the co-culture. The number of transformed cells was substantially lower in the presence of DNase I, indicating that DNA detection occurred via natural transformation in the co-cultures (**Fig. 4B**). Antibiotic resistance is prevalent in microbiomes and may not be used universally as a treatment for the donor cells. Therefore, we tested if heat treatment could be used to efficiently release donor cell DNA. Incubation of *E. coli* at 90°C for 10 minutes substantially enhanced the EC sensor detection limit (5×10^6^ CFU/mL) compared to the addition of spectinomycin (**Fig. 4C**). In sum, detection of target DNA sequences directly from crude samples in the absence of DNA purification could enable the deployment of the DNA sensors for different future applications.

To evaluate the robustness of the DNA sensing function, we characterized the performance of the DNA sensors for multiplexed detection of spike-in bacteria in the presence of cecal contents derived from germ-free mice that were orally gavaged with a defined bacterial consortium (Methods). Mouse ceca contain other bacterial species, host cells and other chemical compounds (e.g. dietary factors), and thus can be used to evaluate the robustness of the DNA sensor. To this end, we introduced varying amounts of *E. coli* and *S. typhimurium* into 10 mg of mouse ceca, incubated these samples at 90°C for 10 minutes, and then transferred the samples into a mixed culture containing the EC-G and ST-R sensors (**Fig. 4D**). Our results demonstrated that both sensors can detect *E. coli* and *S. typhimurium* cells in ceca without DNA extraction. In particular, the EC and ST sensors displayed a detection limit of 10^7^ CFU/mL (**Fig. 4D**). In samples containing a single donor species, high density of *E. coli* or *S. typhimurium* (10^8^ CFU/mL) yielded infrequent false positives for the multiplexed DNA detection. This suggests that further optimization of the DNA sensors may be needed in complex samples containing high donor cell densities that are heat treated (**Fig. S10**). In sum, we show that the cell-based DNA sensors can robustly perform DNA detection of heat-treated samples that contain the target sequence.

## DISCUSSION

Here we engineered the naturally competent bacterium *B. subtilis* to sense and respond to specific DNA sequences. DNA sensing can be achieved for purified DNA or eDNA released from pre-treated samples containing donor cells harboring the target sequence. We demonstrate that DNA sensing can be sensitive and specific, and multiplexed sensing can be achieved by engineering sensors with orthogonal reporter genes. Detection of species using a living cell-based DNA sensor strain opens avenues for future research for versatile sensing of species and does not rely on chemical or physical signals^31,32^. Since our circuit design is modular, customized sensors could be constructed in the future for the detection of sequences derived from diverse organisms including viruses, fungi and mammalian cells.

The DNA detection limit is impacted by the frequency of background mutations and could be improved by reducing the background genetic mutation rate. For example, counter-selectable markers^33^ that can achieve lower background mutation rate could be used to optimize the strength of negative growth selection. The mutation rate can also be reduced by deleting endogenous genes in *B. subtilis* that promote mutagenesis such as the transcription conflict factor *mfd*^34^. The reduction of mutation rates is also critical for long-term implementation of living DNA sensors in the environments^35^.

The time required for DNA detection using the cell-based DNA sensors is relatively slow compared to other diagnostic methods^36^. In particular, the total time required for DNA detection including transformation and selection is approximately one day, which is not suitable for certain applications that require a rapid response. To reduce the detection time, directed evolution or rational design of the natural competence pathway could be used for enhancing the transformation efficiency of *B. subtilis*^37^. This in turn would reduce the time required for transformation and detection of GFP in liquid media. In addition, some naturally competent bacteria such as *Streptococcus pneumoniae* can achieve 50% transformation efficiency^38^. This transformation efficiency is substantially higher than our constructed DNA sensors (10^-5^-10^-4^). With a suitable chassis with a high transformation ability, GFP expression could be observed more rapidly following transformation, which requires a few hours.

The living DNA sensors have potential for *in vitro* DNA detection applications. One unique feature of the cell-based DNA sensor is the long homology within the landing pad region of the circuit, distinct from PCR-based methods that use short recognition sequences. Therefore, the target DNA sequence can be specified at the level of genes or pathways (i.e. biosynthetic gene clusters), and the DNA sensor could be used to mine such sequences from metagenomic DNA^39^. In addition, the access to NGS sequencing or multiplexed qPCR may not be widely available^36,40^. By contrast, the cell-based DNA detection is relatively simple and cost-effective. The DNA sensors may be suitable for large-scale screening with limited experimental resources. Further, *B. subtilis* sensors could be stored as spores for easy and long-term storage^41^. The metrics used in this study (sequence identity similarity and coverage) should be systematically examined using existing sequencing data and experimental characterization to elucidate sequence design rules for homologous recombination. In addition, tools from machine learning could be used to predict the impact of landing pad sequences on the fitness of *B. subtilis* to minimize any negative effects on growth rate^42^.

One of the most unique aspects of this system is the potential for *in situ* DNA detection. *B. subtilis* has been shown to colonize or reside temporarily in diverse environments including soil and the mammalian gastrointestinal tract^43,44^, enabling *in situ* DNA monitoring. For example, living DNA sensors could be introduced into gastrointestinal tract or plant-associated environment to monitor microbiome dynamics by sensing and recording in real-time^45^. The sensing mechanisms could be coupled to the release of antimicrobials to target specific pathogens^46^. A recent study demonstrated that the naturally competent bacterium *Acinetobacter baylyi (A. baylyi*) can be engineered to detect tumor DNA in the mouse colon^47^, demonstrating a potential application of *in situ* DNA detection. In their study, the native CRISPR system in *A. baylyi* was exploited to detect a single mutation in the *KRAS* gene in cancer cells. Similar CRISPR systems could be incorporated into our current circuit design in the future to discriminate between single-nucleotide differences. In sum, we believe that engineering DNA-sensing bacteria could open new avenues for both *in vitro* and *in situ* applications in the future.

## MATERIALS AND METHODS

### Plasmid and strain construction

All DNA sensor strains were derived from *B. subtilis* PY79. Plasmids constructed in this work are listed in **Table S1**. The pAX01-comK plasmid was purchased from Bacillus Genetic Stock Center (BGSC ID: ECE222) to introduce P_xylA_-*comK* at the *lacA* locus in *B. subtilis* PY79 by the selection of MLS (1 μg/mL erythromycin from Sigma-Aldrich and 25 μg/mL lincomycin from Thermo Fisher Scientific) to enhance the transformation efficiency in LB^17,48^. Genes of fluorescent protein GFP(Sp), mCherry, and mTagBFP were cloned from plasmid pDR111_GFP(Sp)^10^ (BGSC ID: ECE278), plasmid mCherry_Bsu^11^ (BGSC ID: ECE756), and plasmid mTagBFP_Bsu^11^ (BGSC ID: ECE745) to construct fluorescent reporter plasmids pOSV00170, pOSV00455 and pOSV00456, respectively. The fluorescent reporter was introduced at the *ycgO* locus by the selection of 5 μg/mL chloramphenicol (MilliporeSigma). The null DNA detection plasmid pOSV00157 was composed of Repressor *lacI* and IPTG-inducible toxin-antitoxin system Phyperspank-*txpA*-*ratA* and can be introduced at the *amyE* locus by the selection of 100 μg/mL spectinomycin (Dot Scientific). The toxin-antitoxin system P_hyperspank_-*txpA-ratA* was PCR amplified from *B. subtilis* 168 gDNA.

*B. subtilis*, *E. coli*, *S. typhimurium*, *S. aureus* and *S. epidermidis* were all cultured at 37°C in Lennox LB medium (MilliporeSigma). *C. difficile* and gut bacterial strains *A. caccae*, *B. thetaiotaomicron*, *C. asparagiforme*, *C. hiranonis*, *and B. longum* were cultured at 37°C in YBHI medium in an anaerobic chamber (Coy Laboratory). YBHI medium is Brain-Heart Infusion Medium (Acumedia Lab) supplemented with 0.5% Bacto Yeast Extract (Thermo Fisher Scientific), 1 mg/mL D-Cellobiose (MilliporeSigma), 1 mg/mL D-maltose (MilliporeSigma), and 0.5 mg/mL L-cysteine (MilliporeSigma). The gDNA of each species was extracted using DNeasy Blood & Tissue Kit (Qiagen). For *S. aureus* gDNA extraction, 0.1 mg/mL Lysostaphin (MilliporeSigma) was added in the pre-treatment step in combination with enzymatic lysis buffer (Qiagen). Bacterial strains are listed in **Table S2**.

The target sequences *xdhABC* were PCR amplified from *E. coli* MG1655 gDNA (NCBI Reference Sequence: NC_000913.3; Location: 3001505-3004004 and 3004005-3006504),*sipBCDA* from *Salmonella enterica* serovar Typhimurium LT2 ATCC 700720 (NCBI Reference Sequence: NC_003197.2; Location: 3025979-3028478 and 3028479-3030978), *hemEH* from *S. aureus* DSM 2569 (GenBank: LHUS02000002.1; Location: 553-2770 and 2864-5638), and *pheST* from *C. difficile* DSM 27147 (GenBank: FN545816.1; Location: 770923-773144 and 773157-775686) to construct a set of plasmids (pOSV00169, pOSV00205, pOSV00206, pOSV00207, pOS00208, pOSV00292, pOSV00459 and pOSV00475) using restriction enzymes BamHI-HF (New England Biolabs) and EcoRI-HF (New England Biolabs) or Golden Gate Assembly Mix (New England Biolabs). DNA sequences of genetic parts are listed in **Table S3**.

All plasmids were constructed using *E. coli* DH5α and transformed into *B. subtilis* using MC medium^49^. MC medium is composed of 10.7 g/L potassium phosphate dibasic (Chem-Impex International), 5.2 g/L potassium phosphate monobasic (MilliporeSigma), 20 g/L glucose (MilliporeSigma), 0.88 g/L sodium citrate dihydrate (MilliporeSigma), 0.022 g/L ferric ammonium citrate (MilliporeSigma), 1 g/L Oxoid casein hydrolysate (Thermo Fisher Scientific), 2.2 g/L potassium L-glutamate (MilliporeSigma), and 20 mM magnesium sulfate (MilliporeSigma). Plasmids were extracted from *E. coli* DH5α using Plasmid Miniprep Kit (Qiagen) for transformation of *B. subtilis. B. subtilis* was inoculated into MC medium and incubated at 37°C for 2 hours. Extracted plasmids were added into B. subtilis cell culture and incubated 37°C for another 4 hours. Transformed *B. subtilis* were selected on the LB agar plate with selective antibiotics. Double crossover was verified for colonies by the replacement of a different antibiotic resistance gene at the integration locus.

### DNA detection using cell-based sensors

DNA sensor strain was inoculated from the −80°C glycerol stock into LB medium with 100 μg/mL spectinomycin and incubated at 37°C with shaking (250 rpm) for 14 hours. On the next day, the OD600 of overnight culture was measured by NanoDrop One (Thermo Fisher Scientific) and diluted to OD0.1 in 1 mL LB in 14 mL Falcon™ Round-Bottom Tube (Thermo Fisher Scientific) supplemented with 50 mM xylose (Thermo Fisher Scientific) and 100 μg/mL spectinomycin (Dot Scientific). Xylose was added to induce the competence. Spectinomycin was added to avoid contamination from other bacteria but it was not required. The sample gDNA was quantified by the Quant-iT dsDNA Assay Kit (Thermo Fisher Scientific) and supplemented in sensor culture with known concentration. The DNA sensor culture was incubated at 37°C with shaking (250 rpm) for 10 hours for transformation. Culture of transformed sensors (5 μL) was plated onto a 12-well plate (Thermo Fisher Scientific) for selection. In these plates, each well contained 1 mL LB agar supplemented with 2 mM IPTG (Bioline), 5 μg/mL chloramphenicol (MilliporeSigma), and MLS (1 μg/mL erythromycin from Sigma-Aldrich and 25 μg/mL lincomycin from Thermo Fisher Scientific). Antibiotics were used in agar to avoid contamination. GFP-expressing colonies were imaged using Azure Imaging System 300 (Azure Biosystems) using Epi Blue LED Light Imaging with 50 millisecond exposure time. CFU was counted manually. Transformation efficiency is defined as the ratio of CFU on selective plates (transformed *B. subtilis* with GFP expression) to the CFU on non-selective plate (total *B. subtilis*). To count CFU of total *B. subtilis*, cell culture was serially diluted in phosphate-buffered saline (PBS) (Dot Scientific) and plated onto LB agar plate supplemented with 5 μg/mL chloramphenicol and MLS. Since the total *B. subtilis* CFU was similar for most conditions, CFU of transformed cells was used to indicate detection efficiency.

To quantify the sensitivity or specificity of sensors, cell culture was transferred to liquid LB medium with a 1:20 dilution after transformation with serially diluted DNA from 1500 ng/mL to 1 ng/mL. To test the specificity towards gDNA from different strains, 100 ng/mL purified DNA was used for transformation. LB medium was supplemented with 2 mM IPTG (Bioline), 5 μg/mL chloramphenicol (MilliporeSigma), and MLS. Diluted cell culture was transferred to a 96-well black and clear-bottom CELLSTAR^®^ microplate with 100 uL volume in each well (Greiner Bio-One). Plate was sealed with Breathe-Easy Adhesive Microplate Seals (Thermo Fisher Scientific) and incubated in the SPARK Multimode Microplate Reader (TECAN) at 37°C with shaking for time-series OD600 and GFP measurements. A threshold of GFP fluorescence 400 was used to determine the detection time for different DNA or different concentrations. The threshold was placed in the region of exponential amplification of GFP across all of the conditions where the difference in the detection time between different conditions was not affected much by the choice of threshold. Unpaired *t*-test was performed to determine if the detection time of specific DNA concentration is different than the background without DNA (N=4). The detection limit is the lowest DNA concentration with a statistical difference. A straight line was fitted to the mean detection time of the four technical replicates with statistical difference versus the logarithmic DNA concentration. DNA mass per mL in detection limit (1 ng, 62.5 ng, 4 ng, and 16 ng) and genome size (4639675 bp, 4857450 bp, 2827820 bp, and 4153430 bp) were converted to chromosome copy number by NEBioCalculator for *E. coli* (2.10×10^5^), *S. typhimurium* (1.25×10^7^), *S. aureus* (1.38×10^6^), and *C. difficile* (3.75×10^6^), respectively.

### Bioinformatic analysis of the specificity of target DNA

To analyze if the target sequence is conserved for the target species, we searched the 5000 bp of *E. coli xdhABC, S. typhimurium sipBCDA, S. aureus hemEH*, and *C. difficile pheST* DNA sequence within the same species in NCBI Nucleotide Collection Database. Nucleotide BLAST was optimized for somewhat similar sequences (blastn). The search was specified to taxid 561 for *E. coli*, taxid 28901 for *S. enterica*, taxid 1280 for *S. aureus*, and taxid 1496 for *C. difficile*. Accession date of data is 2022-08-19. For the same strains with target sequence with more than 95% identity similarity and 95% coverage, the percentage is 99% (N=3462), 94.3% (N=2078), 95.6% (N=1488), and 96.4% (N=139) for *E. coli, S. typhimurium, S. aureus*, and *C. difficile*,respectively. To analyze if the target sequence is conserved in other species, the same search was performed excluding the same species. Homologs with varying identity similarity and coverage were found in different species for the target sequences *E. coli xdhABC* (N=5000), *S. typhimurium sipBCDA* (N=117), *S. aureus hemEH* (N=2993), and *C. difficile pheST* (N=5000). Dot plot of homology coverage and identity similarity of each hit were shown for each target sequence.

### Multiplexed DNA detection using cell-based sensors

Overnight cultures of DNA sensor strains EC-G, ST-R and SA-B were diluted to OD0.1, OD0.1, and OD0.01, respectively, in one single culture containing 1 mL LB supplemented with 50 mM xylose (Thermo Fisher Scientific) and 100 μg/mL spectinomycin (Dot Scientific). Different combinations of gDNA of *E. coli*, *S. typhimurium*, and *S. aureus* (200 ng/mL each) were added into the mixed culture containing three DNA sensor strains in separate 14 mL Falcon tubes (Thermo Fisher Scientific). The DNA sensor strains were incubated at 37°C with shaking (250 rpm) for 10 hours and 5 μL of cell culture was plated onto 12-well plates (Thermo Fisher Scientific). In these plates, each well contained 1 mL LB agar supplemented with 2 mM IPTG, 5 μg/mL chloramphenicol, and MLS. Plates were incubated at 37°C overnight for bacterial growth.

On the next day, each well was imaged using Nikon Eclipse Ti-E Microscope. Brightfield images were collected at 4X magnification using the built-in transilluminator of the microscope. Fluorescence images were collected with the epifluorescence light source X-Cite 120 (Excelitas) and standard band filter cubes including BFP (Excitation: 395/25 nm, Emission: 460/50 nm, Chroma), GFP (Excitation: 470/40 nm, Emission: 525/50 nm, Nikon) and Texas Red (Excitation: 560/40 nm, Emission: 630/70 nm, Nikon) to image BFP, GFP, and RFP, respectively. Pixels were processed with 8×8 binning when taking images. The exposure times for BFP, GFP, RFP were 1.5 ms, 1.5 ms, and 7 ms, respectively. Complete images of each LB agar well were generated from multipoint images after scanning a 24×24 mm area. Once the full images were assembled, each of the four channels were mapped to unity by the minimum and maximum pixel values of all images for that channel using ImageJ. Colonies of different colors were counted manually.

### Multiplexed detection in complex DNA samples using NGS or cell-based DNA sensors

Bacterial species *A. caccae, B. thetaiotaomicron, B. longum, C. asparagiforme, S. typhimurium*,and *S. aureus* were inoculated from the −80°C glycerol stock into YBHI medium and incubated at 37°C anaerobically overnight. On the next day, cell culture of each strain was diluted to OD0.01 in YBHI (Passage 0) and incubated for 24 hours (Passage 1). Cell culture was diluted to OD0.1 into a fresh YBHI and incubated for another 24 hours (Passage 2). At each passage, cell pellet was collected for DNA extraction using DNeasy Blood & Tissue Kit (Qiagen). To extract *S. aureus* gDNA, 0.1 mg/mL Lysostaphin (MilliporeSigma) was added in the pre-treatment step in combination with enzymatic lysis buffer (Qiagen). Purified DNA was stored at −20°C before further processing for NGS or cell-based detection.

To use NGS for characterizing microbial community composition, the V3-V4 region of the 16S rRNA gene was PCR amplified from extracted DNA using custom dual-indexed primers on a 96-well PCR plate (detailed method described in Clark et al. Nat. Comm., 2021^50^). PCR products from each well were pooled and purified using DNA Clean & Concentrator kit (Zymo). The resulting library was sequenced on an Illumina MiSeq using a MiSeq Reagent Kit v2 (500-cycle) to generate 2×250 paired-end reads. Sequencing data were demultiplexed using Basespace Sequencing Hub’s FastQ Generation program. Custom python scripts were used for further data processing and sequences were mapped to the 16S rRNA reference database created using consensus sequences from Sanger sequencing data of monospecies cultures. Relative abundance was calculated as the read count mapped to each species divided by the total number of reads of all species.

To use cell-based DNA sensors to detect *S. typhimurium* and *S. aureus* gDNA in the community DNA, 1000 ng/mL extracted community DNA was added to the mixture of SA-G (OD0.1) and ST-R sensors (OD0.1) for the multiplexed detection of *S. typhimurium* and *S. aureus*.The DNA sensor strains were incubated at 37°C with shaking (250 rpm) for 10 hours and 5 μL of cell culture was plated onto 12-well plates (Thermo Fisher Scientific) containing 1 mL LB agar supplemented with 2 mM IPTG, 5 μg/mL chloramphenicol, and MLS. Plates were incubated at 37°C overnight for bacterial growth. On the next day, each well was imaged using Nikon Eclipse Ti-E Microscope. Brightfield and fluorescence images of GFP and RFP were collected at 4X magnification using the same procedure described in the previous section for the multiplexed DNA detection. Colony numbers were counted manually. The fold change of mean colony numbers compared to Day 1 was compared with the fold change of relative abundance by NGS results.

### Direct detection of target species in the co-culture using cell-based sensors

*E. coli, S. typhimurium*, and *S. aureus* and the corresponding DNA sensor strains were inoculated from the −80°C glycerol stock into LB medium and incubated at 37°C with shaking (250 rpm) for 14 hours. *C. difficile* was separately inoculated in YBHI medium and incubated in an anaerobic chamber (Coy Laboratory). On the next day, cell culture of sensor and target strain were diluted to an OD0.1 each in a single culture containing 1 mL LB supplemented with 50 mM xylose with or without 100 μg/mL spectinomycin in 14 mL Falcon tubes (Thermo Fisher Scientific). Overnight culture of target strains (14 hr) was diluted in PBS (Dot Scientific) and plated onto LB agar plates or YBHI agar plates to determine the initial CFU of target bacteria in the co-culture (OD0.1), which was 1.22×10^8^ CFU/mL, 1.07×10^8^ CFU/mL, 3.2×10^8^ CFU/mL, and 1.1×10^7^ CFU/mL for *E. coli, S. typhimurium*, *S. aureus*, *and C. difficile*, respectively. Sensor and target strains were co-cultured at 37°C with shaking (250 rpm) for 10 hours, and 5 μL of cell culture was plated onto 12-well plates (Thermo Fisher Scientific). Each well contained 1 mL LB agar supplemented with 2 mM IPTG, 5 μg/mL chloramphenicol, and MLS for the selection of transformed *B. subtilis*. The 12-well plates were incubated overnight at 37°C. On the next day, fluorescent colonies were imaged by Azure Imaging System 300 (Azure Biosystems) using the Epi Blue LED Light Imaging with 50 millisecond exposure time. Colonies with GFP expression were counted manually. To determine if the detection of target bacteria was via transformation, 1 μL (1 unit) DNase I (Thermo Fisher Scientific) was added to the 1 mL co-culture. One unit of DNase I can completely degrade 1 μg of plasmid DNA in 10 min at 37°C according to the manufacturer’s specification.

To improve the detection efficiency, overnight culture of *E. coli* was incubated at 90°C in digital dry baths/block heaters (Thermo Fisher Scientific) for 10 min and placed on ice for 3 min before being transferred to the sensor culture containing 1 mL LB, 50 mM xylose, and OD0.1 of sensor strain for detection. Spectinomycin was not used in the sensor culture for heat-treated samples. To test the multiplexed detection of *E. coli* and *S. typhimurium* in mice cecal samples, 10 mg cecal samples were first resuspended with 100 μL LB and 50 mM xylose in 1.7 mL Eppendorf tubes (Dot Scientific). Different amounts of overnight culture of *E. coli* and *S. typhimurium* were spiked into cecal samples. The cecal samples were then incubated at 90°C for 10 min and sat on ice for 3 min before being transferred to the mixed culture of EC-G and ST-R sensors (1 mL LB, 50 mM xylose, OD0.1 of EC-G, and OD0.1 of ST-R) for multiplexed detection. Cecal samples were collected from germ-free mouse experiments following protocols approved by the University of Wisconsin-Madison Animal Care and Use Committee. Briefly, 8-week-old C57BL/6 gnotobiotic male mice (wild-type) were inoculated with 8 human gut bacteria - *Dorea formicigenerans*, *Coprococcus comes*, *Anaerostipes caccae*, *Bifidobacterium longum*, *Bifidobacterium adolescentis, Bacteroides vulgatus, Bacteroides caccae*, and *Bacteroides thetaiotaomicron* via oral gavage. After four weeks of colonization, mice were euthanized for cecal sample collection. Cecal samples were stored at −80°C and thawed for the use in multiplexed detection of spike-in *E. coli* and *S. typhimurium*.

## ACKNOWLEDGEMENTS

We would like to thank Dr. Scott Coyle and Zhejing Xu for the use of Nikon Eclipse Ti2-E Microscope. We also want to thank Yiyi Liu for providing the mice cecal samples. This work was supported by the Defense Advanced Research Projects Agency (DARPA) Grant HR0011-19-2-0002.

## AUTHOR CONTRIBUTIONS

Y.Y.C., B.M.B. and O.S.V. conceived the research. Y.Y.C., and Z.C. performed the experiments. Y.Y.C., Z.C., and X.C. constructed the sensors. T.D.R. processed the microscopic images. T.G.F. assisted with the strain construction. Y.Y.C. and O.S.V. wrote the manuscript. O.S.V. secured the funding.

