## Supplementary Information for "Programming bacteria for multiplexed DNA detection"

**This file includes:**

Figure S1. Plasmid maps for the construction of DNA-sensing *B. subtilis*.
Figure S2. Sequencing of transformed *E. coli* sensor and escape mutants.
Figure S3. Nucleotide BLAST search of homology sequences in EC sensor and ST sensor.
Figure S4. Time-series measurements of GFP expression of ST, SA, CD sensors in liquid medium after transformation.
Figure S5. Time-series OD measurements of DNA sensors in liquid medium after transformation.
Figure S6. Orthogonality test of the four constructed DNA sensors.
Figure S7. Detection efficiency based on number of transformed cells.
Figure S8. Representative fluorescence images of transformed EC-G, ST-R, and SA-B sensors for multiplexed detection.
Figure S9. Negative control for the multiplexed detection in complex DNA samples.
Figure S10. Detection of *E. coli* or *S. typhimurium* in cecal samples using mixed EC-G and ST-R sensors.
Table S1. List of plasmids.
Table S3. List of bacterial strains.
Table S3. Sequences of genetic parts.

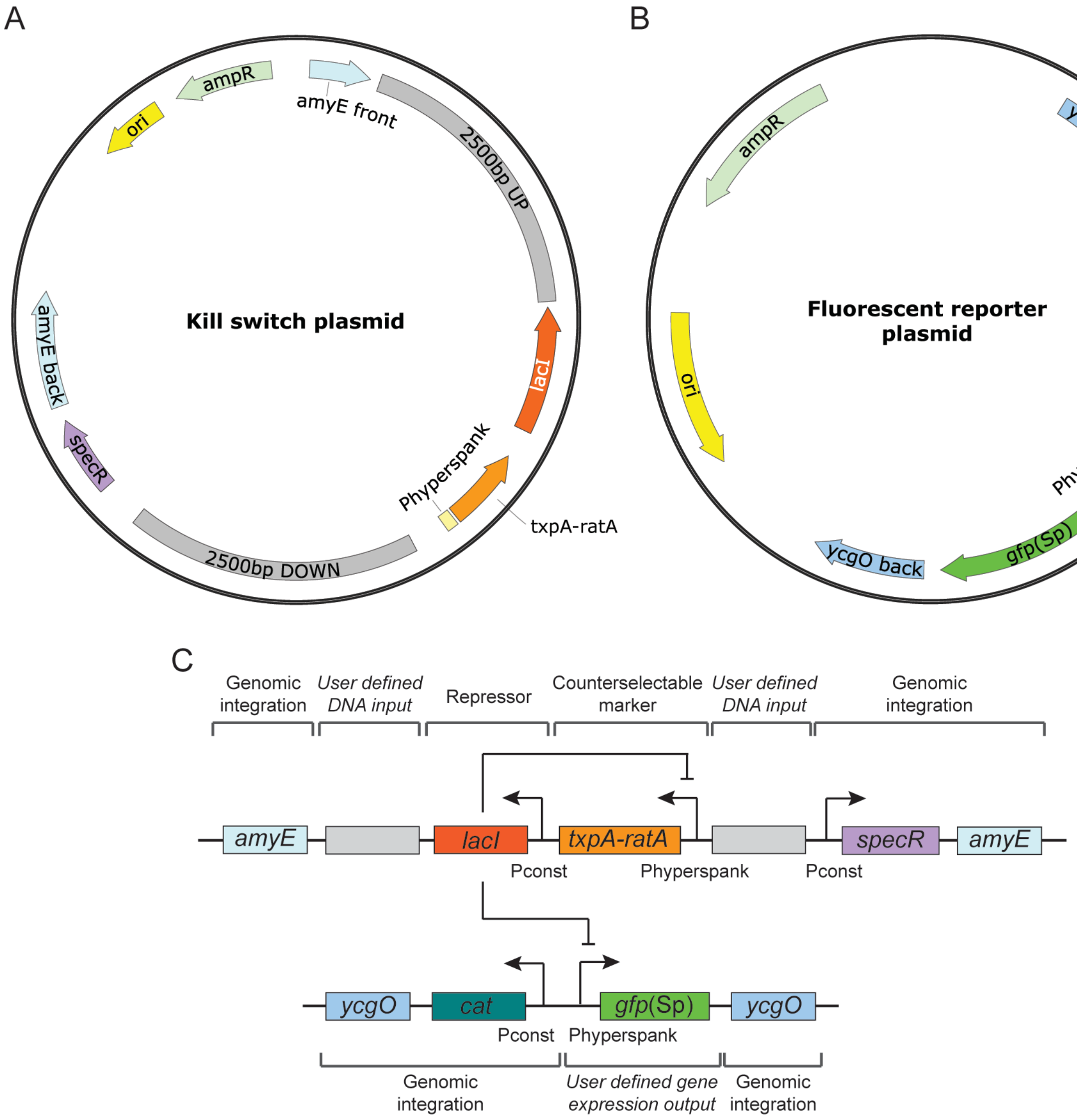


**Figure S1. Plasmid maps for the construction of DNA-sensing *B. subtilis*. (A)** Plasmid constructed for the DNA detection via homologous recombination. The toxin-antitoxin system *txpA-ratA* is regulated by the repressor *lacI*, both of which are flanked by 2.5 kbp target DNA sequence on each side. The plasmid was integrated into the *amyE* locus on the *B. subtilis* PY79 genome by spectinomycin selection. **(B)** Plasmid constructed for the fluorescent reporter after DNA detection. The green fluorescent protein *gfp*(Sp) is regulated by the distal repressor *lacI*. The plasmid was integrated into the *ycgO* locus on *B. subtilis* PY79 genome by chloramphenicol selection. The green fluorescent protein *gfp*(Sp) was codon-optimized for *Streptococcus pneumoniae* and displayed high fluorescence signal in *B. subtilis*[1]. **(C)** Modularity of the synthetic genetic circuit allows customized target DNA sequence as input and gene expression as output.

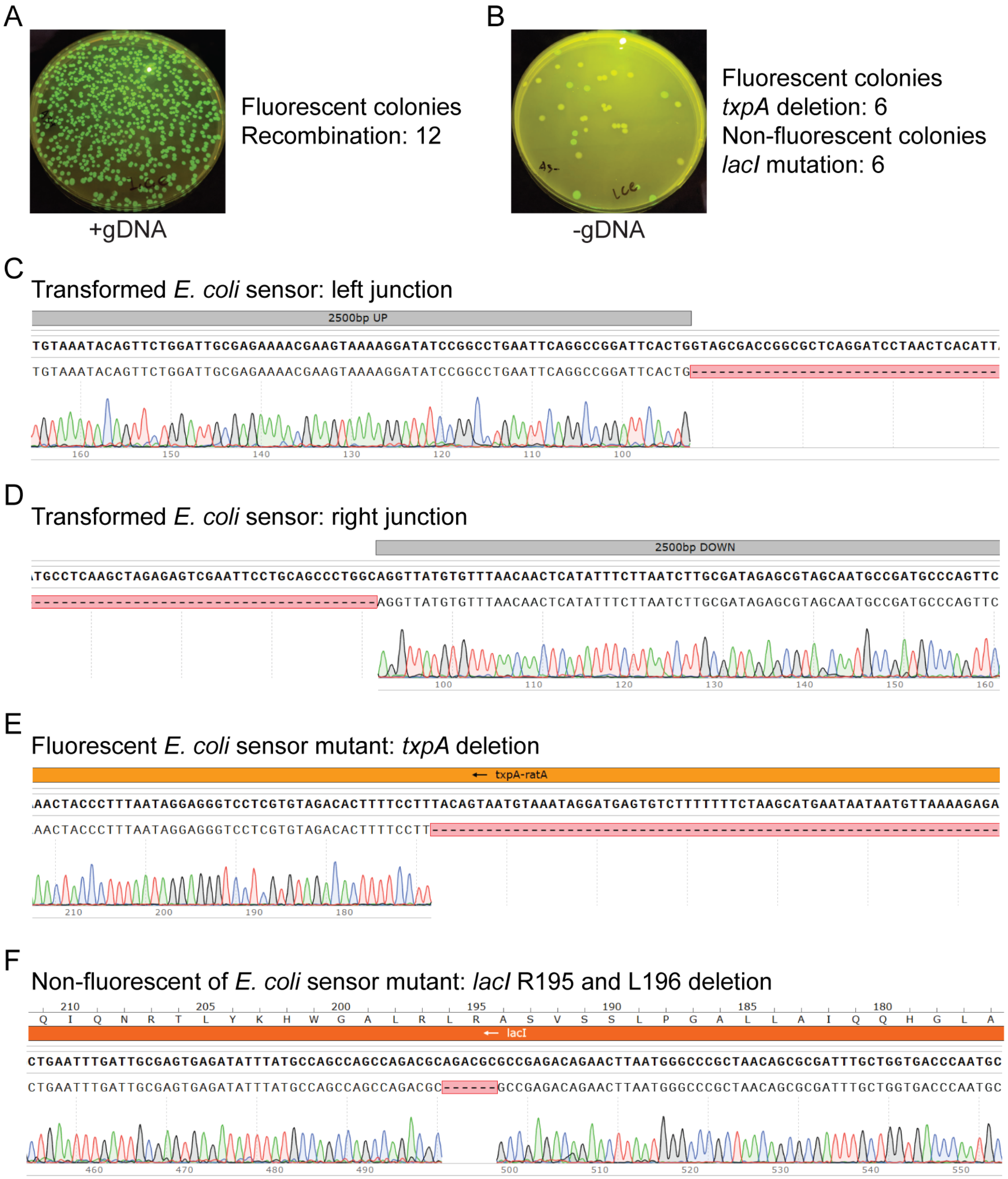


**Figure S2. Sequencing of transformed *E. coli* sensor and escape mutants.** Representative fluorescence images of colonies of *E. coli* DNA sensor on selective agar plate after the transformation of **(A)** 100 ng/mL purified *E. coli* gDNA or **(B)** no DNA. DNA of the circuit in *B. subtilis* genome was PCR amplified from different colonies and sequenced to confirm the homologous recombination for the transformed sensor (N=12) or mutations for the escape mutants (N=12). **(C)(D)** Sanger sequencing of transformed *E. coli* sensor confirmed the homologous recombination at the predicted region that removed the whole cassette of *txpA-ratA* and *lacI*. **(E)** Sanger sequencing of a GFP-expressing escape mutant shows deletion in the toxin *txpA*. The mutant can grow and express GFP in the presence of IPTG due to the non-functional toxin. **(F)** Sanger sequencing of a non-fluorescent escape mutant shows deletion of R195 and

L196 in the repressor *lacI*, which has been shown to affect IPTG binding[2]. The mutant can grow but cannot express GFP in the presence of IPTG due to the non-functional LacI.

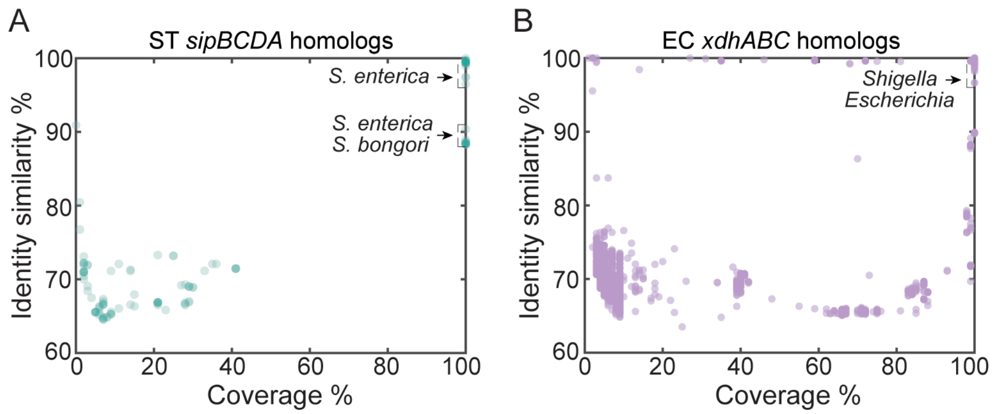


**Figure S3. Nucleotide BLAST search of homology sequences in EC sensor and ST sensor. (A)** Nucleotide BLAST search of 5000 bp *S. typhimurium sipBCDA* in the NCBI database. Each circle represents a homolog found in species other than *S. typhimurium* and its coverage and identity similarity. Homologs were found mostly in the *Salmonella enterica* (*S. enterica*) species but were rarely found in other species. **(B)** Nucleotide BLAST search of 5000 bp *E. coli* MG1655 *xdhABC* in the NCBI database. Each circle represents a homolog found in species other than *E. coli* and its coverage and identity similarity. Homologs were found in closely related *Shigella* and *Escherichia* species. Highly similar homologs were found in the closely related *Shigella* species.

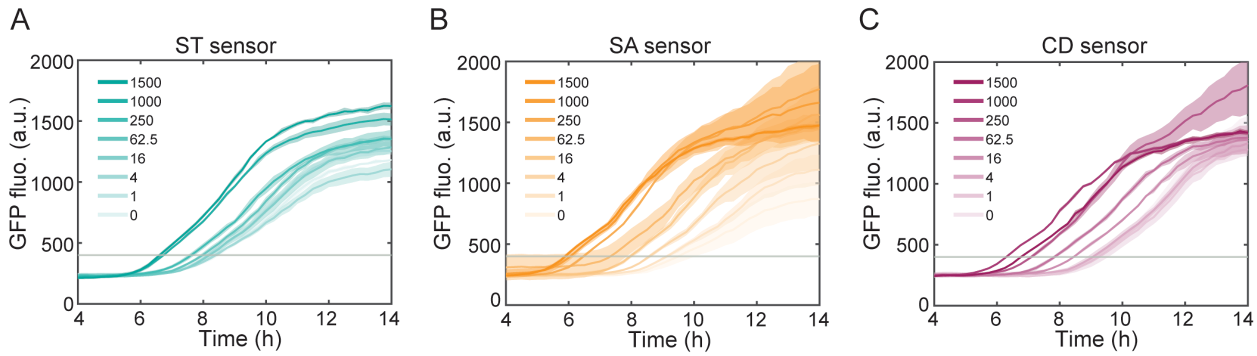


**Figure S4. Time-series measurements of GFP expression of ST, SA, CD sensors in liquid medium after transformation.** Time-series measurements of GFP expression of **(A)** ST sensor, **(B)** SA sensor, and **(C)** CD sensor in liquid medium after the transformation of varying target gDNA concentrations (ng/mL). A threshold of GFP 400 was used to determine the detection time for each gDNA concentration. Line is the average of four technical replicates and the shaded region represents one standard deviation from the average.

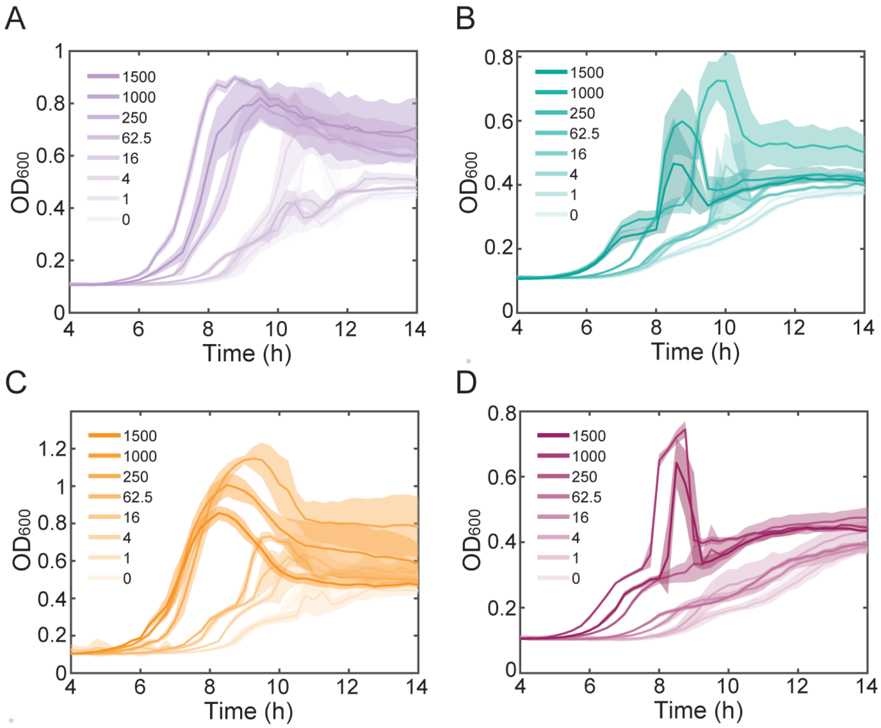


**Figure S5. Time-series OD measurements of DNA sensors in liquid medium after transformation.** Time-series measurements of OD600 absorbance of **(A)** EC sensor, **(B)** ST sensor, **(C)** SA sensor, and **(D)** CD sensor in liquid medium after the transformation of varying target gDNA concentrations (ng/mL). Line is the average of four technical replicates and the shaded region represents one standard deviation from the average. Cell growth correlated with DNA concentration during transformation.

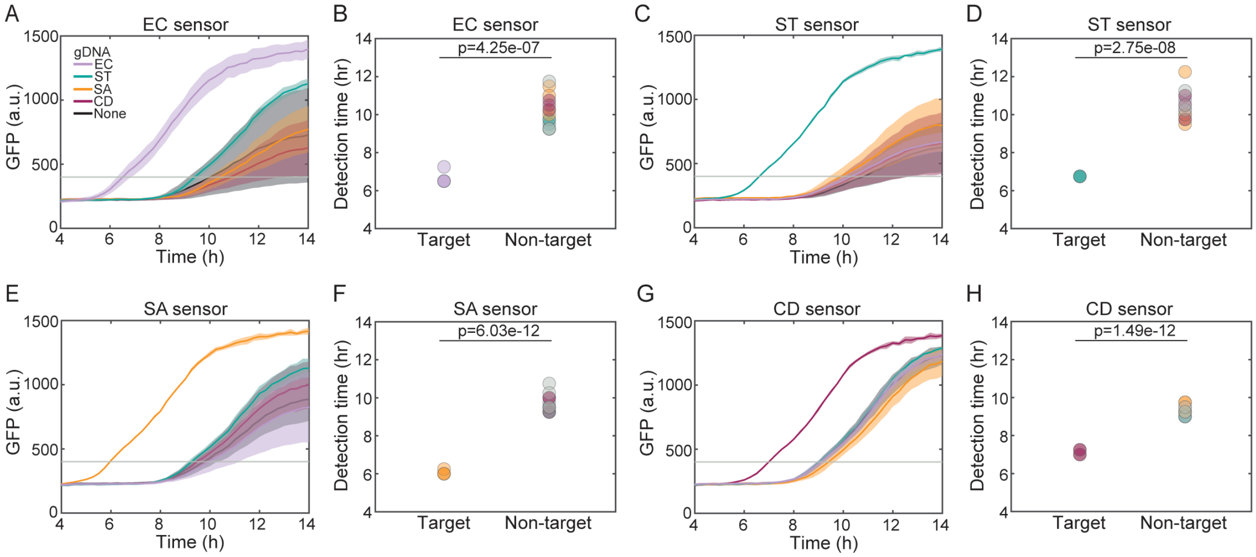


**Figure S6. Orthogonality test of the four constructed DNA sensors.** Time-series measurements of GFP expression of **(A)** EC sensor, **(C)** ST sensor, **(E)** SA sensor, and **(G)** CD sensor in liquid medium after the transformation of gDNA extracted from different strains or no gDNA. Line is the average of four technical replicates and the shaded region represents one standard deviation from the average. A threshold of GFP 400 was used to determine the detection time for the target gDNA and non-target gDNA or no gDNA for **(B)** EC sensor, **(D)** ST sensor, **(F)** SA sensor, and **(H)** CD sensor. Unpaired *t*-test was performed to determine if the detection time for the target gDNA is different from non-target gDNA or no gDNA. Based on the calculated *p*-values, sensors expressed GFP hours earlier only in the presence of target gDNA.

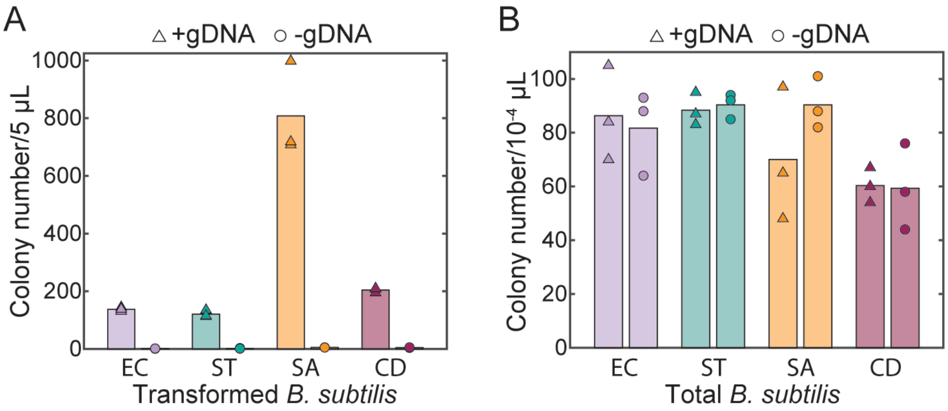


**Figure S7. Detection efficiency based on number of transformed cells. (A)** Colony number per 5 µL for transformed *B. subtilis* with or without 100 ng/mL target gDNA. **(B)** Colony number per $10^{-4}$ µL for total *B. subtilis* with or without 100 ng/mL target gDNA. Bar represents the average of three biological replicates. The transformation efficiency in **Figure 2A** was calculated by the ratio of the density of transformed *B. subtilis* to the density of total *B. subtilis* shown here.

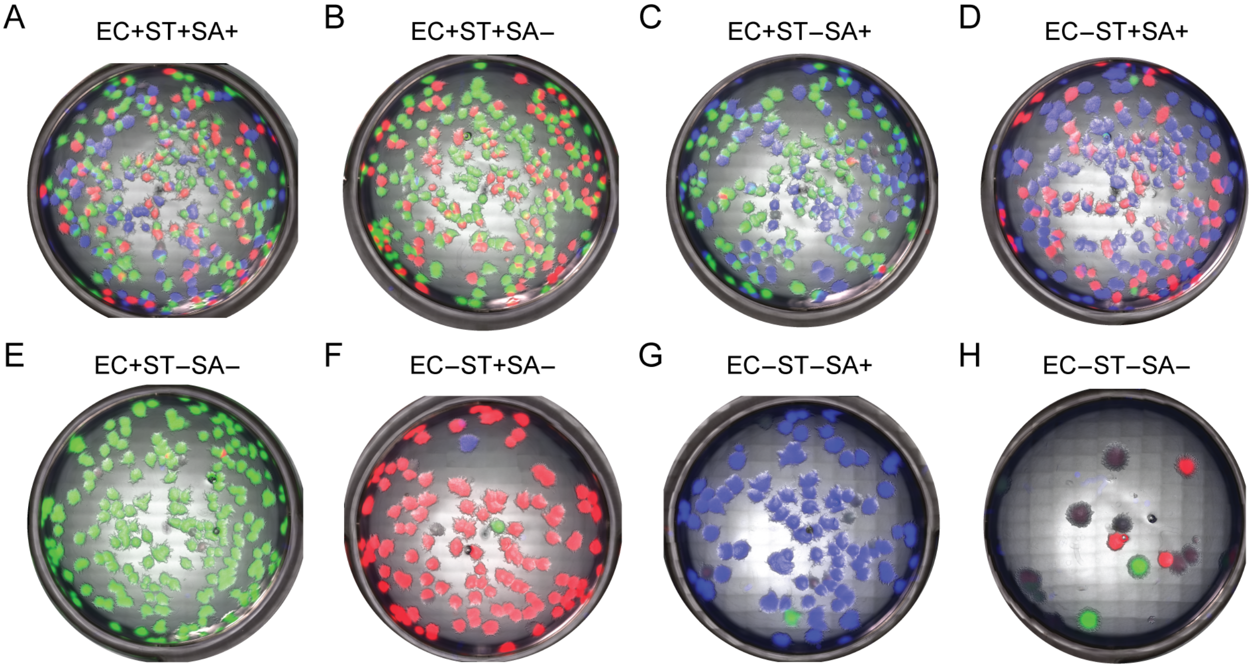

**Figure S8. Representative fluorescence images of transformed EC-G, ST-R, and SA-B sensors for multiplexed detection.** Colonies with different fluorescence on agar plate after the transformation of combinations of extracted gDNA in a mixture of EC-G, ST-R, and SA-G sensors. Sensors were transformed with **(A)** *E. coli*, *S. aureus* and *S. typhimurium* gDNA, **(B)** *E. coli* and *S. typhimurium* gDNA, **(C)** *E. coli* and *S. aureus* gDNA, **(D)** *S. typhimurium* and *S. aureus* gDNA, **(E)** *E. coli* gDNA, **(F)** *S. typhimurium* gDNA, **(G)** *S. aureus* gDNA, and **(H)** no gDNA.

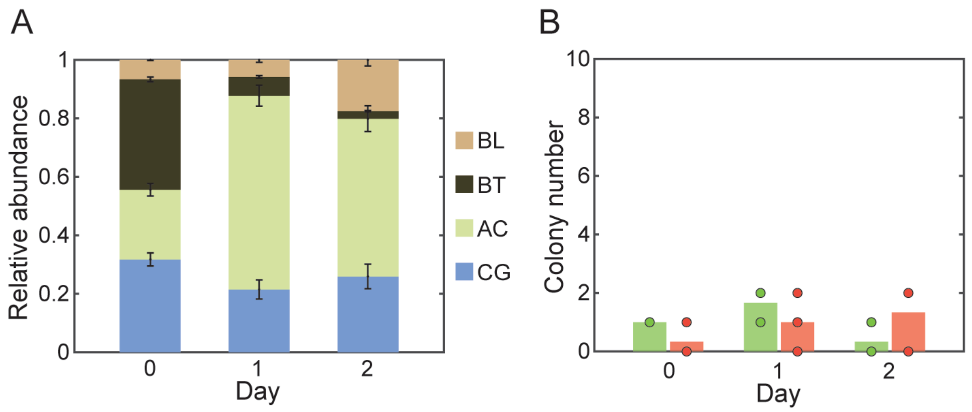


**Figure S9. Negative control for the multiplexed detection in complex DNA samples. (A)** Relative abundance of six species in a synthetic gut microbial community composed of *Bifidobacterium longum* (BL), *Bacteroides thetaiotaomicron* (BT), *Anaerostipes caccae* (AC), and *Clostridium asparagiforme* (CG). The four bacteria were co-cultured in liquid medium anaerobically for 24 hours and cell culture was diluted in fresh medium once for species to continue their competition. 16S rRNA gene of each strain was PCR amplified and sequenced using NGS to determine the relative abundance over time. **(B)** Numbers of GFP or RFP-expressing colonies on agar plate after the transformation of gDNA extracted from the community at different time in a mixture of SA-G and ST-R sensors. Bar represents the average of three technical replicates of 16S rRNA sequencing. Only a few colonies appeared on agar plates, indicating that sensors did not detect the gDNA of *S. aureus* or *S. typhimurium*. Bar represents the average of three technical replicates.

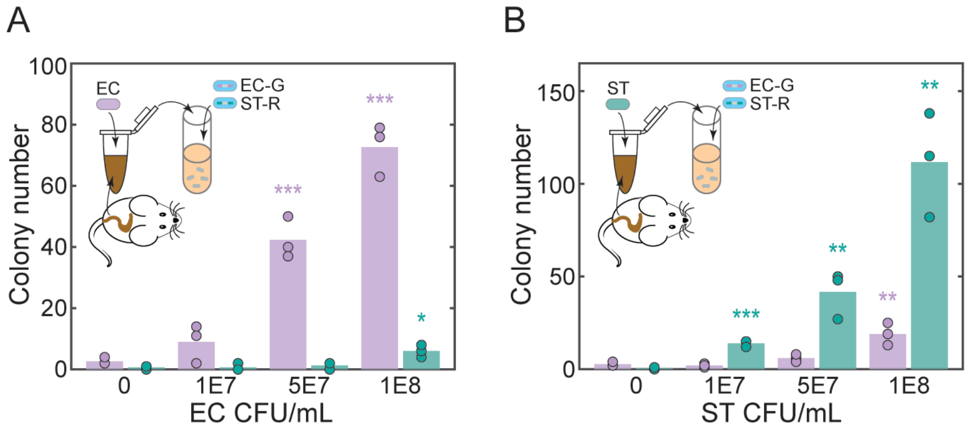


**Figure S10. Detection of *E. coli* or *S. typhimurium* in cecal samples using mixed EC-G and ST-R sensors.** Colony number of transformed EC-G and ST-R sensors co-cultured with heattreated cecal samples spiked in with different amounts of **(A)** *E. coli* or **(B)** *S. typhimurium*. High density ($10^8$ CFU/mL) of *E. coli* and *S. typhimurium* can lead to false positive results for ST-R and EC-G, respectively. Unpaired *t*-test was performed to determine if the colony number is different from no cell condition, and *, **, and *** denote *p*-values < 0.05, 0.01, and 0.001, respectively. Bar represents the average of three technical replicates.

**Table S1. List of plasmids.**

| Plasmid | Description | Genotype |
|---|---|---|
| pAX01-comK[3] | Xylose-inducible *comK* | *lacA*(up), *erm*, $P_{xylA}$-*comK*, *xylR*, *lacA*(down) |
| pOSV00170 | GFP reporter | *ycgO*(up), *cat*, $P_{hyperspank}$-*gfp*, *ycgO*(down) |
| pOSV00455 | RFP reporter | *ycgO*(up), *cat*, $P_{hyperspank}$-*rfp*, *ycgO*(down) |
| pOSV00456 | BFP reporter | *ycgO*(up), *cat*, $P_{hyperspank}$-*bfp*, *ycgO*(down) |
| pOSV00157 | Detection plasmid without target sequence | *amyE*(up), *lacI*, $P_{hyperspank}$-*txpA-ratA*, *spec*, *amyE*(down) |
| pOSV00169 | Detection plasmid with 500 bp EC homology | *amyE*(up), 0.5 kbp EC(up), *lacI*, $P_{hyperspank}$-*txpA-ratA*, 0.5 kbp EC(down), *spec*, *amyE*(down) |
| pOSV00205 | Detection plasmid with 1000 bp EC homology | *amyE*(up), 1 kbp EC(up), *lacI*, $P_{hyperspank}$-*txpA-ratA*, 1 kbp EC(down), *spec*, *amyE*(down) |
| pOSV00206 | Detection plasmid with 1500 bp EC homology | *amyE*(up), 1.5 kbp EC(up), *lacI*, $P_{hyperspank}$-*txpA-ratA*, 1.5 kbp EC(down), *spec*, *amyE*(down) |
| pOSV00207 | Detection plasmid with 2000 bp EC homology | *amyE*(up), 2 kbp EC(up), *lacI*, $P_{hyperspank}$-*txpA-ratA*, 2 kbp EC(down), *spec*, *amyE*(down) |
| pOSV00208 | Detection plasmid with 2500 bp EC homology | *amyE*(up), 2.5 kbp EC(up), *lacI*, $P_{hyperspank}$-*txpA-ratA*, 2.5 kbp EC(down), *spec*, *amyE*(down) |
| pOSV00292 | Detection plasmid with 2500 bp ST homology | *amyE*(up), 2.5 kbp ST(up), *lacI*, $P_{hyperspank}$-*txpA-ratA*, 2.5 kbp ST(down), *spec*, *amyE*(down) |
| pOSV00459 | Detection plasmid with 2500 bp SA homology | *amyE*(up), 2.5 kbp SA(up), *lacI*, $P_{hyperspank}$-*txpA-ratA*, 2.5 kbp SA(down), *spec*, *amyE*(down) |
| pOSV00475 | Detection plasmid with 2500 bp CD homology | *amyE*(up), 2.5 kbp CD(up), *lacI*, $P_{hyperspank}$-*txpA-ratA*, 2.5 kbp CD(down), *spec*, *amyE*(down) |

**Table S2. List of bacterial strains.**

| Strain | Description | Genotype |
|---|---|---|
| msOSV00487 | EC-sensor with 500 bp homology | *B. subtilis* PY79 *amyE*::0.5 kbp EC(up), *lacI*, $P_{hyperspank}$-*txpA-ratA*, 0.5 kbp EC(down), *spec*; *ycgO*::*cat*, $P_{hyperspank}$-*gfp*; *lacA*::*erm*, $P_{xylA}$-*comK*, *xylR* |
| mOSV00580 | EC-sensor with 1000 bp homology | *B. subtilis* PY79 *amyE*::1 kbp EC(up), *lacI*, $P_{hyperspank}$-*txpA-ratA*, 1 kbp EC(down), *spec*; *ycgO*::*cat*, $P_{hyperspank}$-*gfp*; *lacA*::*erm*, $P_{xylA}$-*comK*, *xylR* |
| mOSV00581 | EC-sensor with 1500 bp homology | *B. subtilis* PY79 *amyE*::1.5 kbp EC(up), *lacI*, $P_{hyperspank}$-*txpA-ratA*, 1.5 kbp EC(down), *spec*; *ycgO*::*cat*, $P_{hyperspank}$-*gfp*; *lacA*::*erm*, $P_{xylA}$-*comK*, *xylR* |

| | | |
|---|---|---|
| mOSV00582 | EC-sensor with 2000 bp homology | *B. subtilis* PY79 *amyE*::2 kbp EC(up), *lacI*, $P_{hyperspank}$-*txpA-ratA*, 2 kbp EC(down), *spec*; *ycgO*::*cat*, $P_{hyperspank}$-*gfp*; *lacA*::*erm*, $P_{xylA}$-*comK*, *xylR* |
| msOSV00495 | EC-sensor (EC-G sensor) with 2500 bp homology | *B. subtilis* PY79 *amyE*::2.5 kbp EC(up), *lacI*, $P_{hyperspank}$-*txpA-ratA*, 2.5 kbp EC(down), *spec*; *ycgO*::*cat*, $P_{hyperspank}$-*gfp*; *lacA*::*erm*, $P_{xylA}$-*comK*, *xylR* |
| msOSV00605 | ST sensor | *B. subtilis* PY79 *amyE*::2.5 kbp ST(up), *lacI*, $P_{hyperspank}$-*txpA-ratA*, 2.5 kbp ST(down), *spec*; *ycgO*::*cat*, $P_{hyperspank}$-*gfp*; *lacA*::*erm*, $P_{xylA}$-*comK*, *xylR* |
| msOSV00906 | SA sensor | *B. subtilis* PY79 *amyE*::2.5 kbp SA(up), *lacI*, $P_{hyperspank}$-*txpA-ratA*, 2.5 kbp SA(down), *spec*; *ycgO*::*cat*, $P_{hyperspank}$-*gfp*; *lacA*::*erm*, $P_{xylA}$-*comK*, *xylR* |
| msOSV01005 | CD sensor | *B. subtilis* PY79 *amyE*::2.5 kbp CD(up), *lacI*, $P_{hyperspank}$-*txpA-ratA*, 2.5 kbp CD(down), *spec*; *ycgO*::*cat*, $P_{hyperspank}$-*gfp*; *lacA*::*erm*, $P_{xylA}$-*comK*, *xylR* |
| msOSV01009 | ST-R sensor | *B. subtilis* PY79 *amyE*::2.5 kbp ST(up), *lacI*, $P_{hyperspank}$-*txpA-ratA*, 2.5 kbp ST(down), *spec*; *ycgO*::*cat*, $P_{hyperspank}$-*rfp*; *lacA*::*erm*, $P_{xylA}$-*comK*, *xylR* |
| msOSV01008 | SA-B sensor | *B. subtilis* PY79 *amyE*::2.5 kbp SA(up), *lacI*, $P_{hyperspank}$-*txpA-ratA*, 2.5 kbp SA(down), *spec*; *ycgO*::*cat*, $P_{hyperspank}$-*bfp*; *lacA*::*erm*, $P_{xylA}$-*comK*, *xylR* |
| usOSV00264 | *Escherichia coli* MG1655 | |
| usOSV00197 | *Salmonella enterica* serovar Typhimurium LT2 ATCC 700720 | |
| usOSV00113 | *Staphylococcus aureus* DSM 2569 | |
| usOSV00095 | *Clostridium difficile* DSM 27147 | |
| usOSV00165 | *Staphylococcus epidermidis* ATCC 14990 | |
| usOSV00046 | *Clostridium hiranonis* DSM 13275 | |
| usOSV00157 | *Anaerostipes caccae* DSMZ 14662 | |
| usOSV00011 | *Bacteroides thetaiotaomicron* ATCC 29148 | |
| usOSV00041 | *Clostridium asparagiforme* DSM 15981 | |
| usOSV00067 | *Bifidobacterium longum* subs. *infantis* DSM 20088 | |

**Table S3. Sequences of genetic parts.**

| Part | Sequence |
|---|---|
| $P_{hyperspank}$-*txpA*-*ratA* | ctcgagggtaaatgtgagcactcacaattcattttgcaaaagttgttgactttatctacaaggtgtggcataatgtgtgtaattgtgagcggataacaattaagcttacataaggaggaactactATGTCGACCTATGAATCTCTAATGGTCATGATCGGCTTTGCCAATTTAATAGGCGGGATTATGACATGGGTAATATCTCTTTTAACATTATTATTCATGCTTAGAAAAAAAGACACTCATCCTATTTACATTACTGTAAAGGAAAAGTGTCTACACGAGGACCCTCCTATTAAAGGGTAGTTTCTTTTTTAAAAGCTAGAGTGCTGCCACACTCTGGCTTTTATATTTTAGCATTTCTCATGAAAGTAACACACATTAACAAGTGGTAATGTGGTAATGTGGTACCAACTATAAGCTTACGCCAGTAGTTGCAATACTTTTGCTTGGCACCATTATAACATGAATATATATTGATTATATAATTATTTGTATCTTTTATTTGTTACTTTTTTTATCTATGAGTTCAAAATGACCTGATCATAGAAGCCTTAACCCTTTTTCTTTTATTAAAAACCCTCGGATTATGAAAGTGTTATGGTACAATATGGTTTAGTATAAATGAATATTGGCTTTCAACATCTCAAGGGCGGTCTGGCTCACTCCCTCATGAAAGGGGGTGATGCACGTGTCAACATTTCAAGCATTAATGCTTATGCTTGCTTTCGGGTCATTTATAATTGCCCTGTTGACTTATATAAAGAAGAAATAGACCCACCCCTTGAGCTCGGCAAAGTAAAAGGGTAA |
| *gfp*(*Sp*) | ATGGTTTCTAAAGGTGAAGAATTGTTTACAGGTGTTGTTCCAATTTTGGTTGAATTGGATGGTGATGTTAATGGTCATAAATTTTCTGTTTCTGGTGAAGGTGAAGGTGATGCTACATACGGTAAATTGACATTGAAATTTATTTGTACAACTGGTAAATTGCCAGTTCCTTGGCCAACATTGGTTACAACATTTGCTTATGGTTTGCAATGTTTTGCTCGTTATCCAGATCACATGAAACAACATGATTTCTTTAAATCTGCTATGCCAGAAGGTTATGTTCAAGAACGTACAATCTTTTTCAAGGATGATGGTAATTATAAGACACGTGCTGAGGTTAAGTTTGAAGGTGATACATTGGTTAATCGTATCGAATTGAAGGGTATCGATTTTAAAGAAGATGGTAATATCTTGGGTCATAAATTGGAATATAATTATAATTCTCATAATGTTTATATCATGGCTGATAAACAAAAGAACGGTATTAAAGTTAATTTTAAAATTCGTCATAATATTGAAGATGGTTCTGTTCAATTGGCTGATCATTATCAACAAAATACACCAATTGGTGATGGTCCAGTTTTGTTGCCAGATAATCATTATTTGTCTACACAATCTAAATTGTCTAAAGATCCAAATGAAAAACGTGATCACATGGTTTTGTTGGAATTTGTTACAGCTGCTGGTATTACACATGGTATGGATGAATTGTATAAATAA |
| *rmCherry* | ATGGTTAGCAAAGGCGAAGAGGATAATATGGCGATCATCAAAGAATTTATGCGCTTTAAAGTTCATATGGAAGGCAGCGTTAATGGCCACGAATTTGAAATTGAAGGCGAAGGTGAAGGCAGACCGTATGAAGGCACACAAACAGCAAAACTGAAAGTTACAAAAGGCGGACCGCTGCCGTTTGCATGGGATATTCTGTCACCGCAATTTATGTATGGCAGCAAAGCATATGTTAAACATCCGGCAGATATCCCGGATTATCTGAAACTGTCATTTCCGGAAGGCTTTAAATGGGAACGCGTCATGAATTTTGAAGATGGCGGAGTTGTTACAGTCACACAAGATTCATCACTGCAAGATGGCGAATTTATCTATAAAGTCAAACTGCGTGGCACGAACTTTCCGTCAGATGGCCCTGTTATGCAGAAAAAAACAATGGGCTGGGAAGCATCAAGCGAAAGAATGTATCCGGAAGATGGTGCACTGAAAGGCGAAATTAAACAACGCCTGAAACTTAAAGACGGTGGACATTATGATGCGGAAGTCAAAACAACGTATAAAGCGAAAAAACCTGTTCAACTGCCTGGCGCATATAACGTTAACATTAAACTGGATATCACGAGCCATAACGAAGATTATACAATCGTCGAACAGTATGAAAGAGCAGAAGGACGCCATTCAACAGGCGGAATGGATGAACTGTATAAATACTAG |

| | |
|---|---|
| mTagBFP | ATGAGCGAACTGATCAAAGAAAACATGCATATGAAACTGTACATGGAAG<br>GCACAGTCGATAACCATCACTTTAAATGCACATCAGAAGGCGAAGGCA<br>AACCGTATGAAGGCACACAAACAATGAGAATCAAAGTTGTTGAAGGCG<br>GACCGCTGCCGTTTGCATTTGATATTCTGGCAACATCATTTCTGTATGG<br>CAGCAAAACGTTTATCAATCATACACAAGGCATCCCGGATTTTTTTAAA<br>CAATCATTTCCGGAAGGCTTTACATGGGAACGCGTTACAACATATGAAG<br>ATGGCGGAGTTCTGACAGCAACACAAGATACATCATTGCAAGATGGCT<br>GCCTGATCTATAATGTCAAAATTAGAGGCGTCAACTTTACAAGCAATGG<br>CCCTGTTATGCAGAAAAAAACACTGGGCTGGGAAGCATTTACAGAAAC<br>ACTGTATCCGGCTGATGGCGGACTGGAAGGCAGAAACGATATGGCAC<br>TGAAACTGGTTGGCGGATCACATCTGATTGCAAACATCAAAACAACGTA<br>CCGCTCAAAAAAACCGGCAAAAAATCTGAAAATGCCTGGCGTCTATTAT<br>GTCGATTATAGACTGGAACGCATCAAAGAAGCGAACAACGAAACATAT<br>GTCGAACAACATGAAGTTGCAGTTGCGAGATATTGCGATCTGCCGTCA<br>AAACTGGGCCATAAACTGAATTACTAG |
| EC(up) 0.5 kbp | caattaccatcgaatgcaccattaacgggatgccttttcagcttcacgccgcaccaggcacgccgctctc<br>ggaattactccgcgaacaaggactgctaagtgtcaaacaagggtgctgcgtgggtgaatgtggtgcctgt<br>acggtgttggtcgacggcacagcaatagacagttgcttataccttgccgcctgggctgaaggaaaagag<br>atccgcacgctggaaggtgaagcgaaaggcggaaaactttctcatgttcagcaggcttatgcgaaatcc<br>ggcgcagtgcagtgcgggttttgtacgcctggcctgattatggctaccacggcaatgctggcgaaaccac<br>gcgagaagccattaaccattacggaaattcgtcgcggactggcgggaaatctttgtcgctgcacggggt<br>atcagatgattgtaaatacagttctggattgcgagaaaacgaagtaa |
| EC(down) 0.5 kbp | ttatgtgtttaacaactcatatttcttaatcttgcgatagagcgtagcaatgccgatgcccagttcatcagcaa<br>cttgcttcttgctgttatgacgtgaaagcgcctcgcggatcatttgcttttccatctcctccagcgccgtgccgc<br>ccgcatcatcgagtgacaggtgcgcctcactgacctctgttacatcactttgctccgttgtgccattattcagc<br>agatttggcggcaatagcgtgctgtcgataacttcacctgaaggaaccacgttaaccagatattccatca<br>aattgcttaactcgcgcaggtttccgggccaacgatgcttacgcaatatttcgacgacatcgggagcaatg<br>ccaggataaaccgatcccagacgacgggtatgcagatgtaaaaagtaatgcaccaatagttcaatatct<br>tcctgacgttcacgcagcggtggcagagttatcgggataacattaagtcggtagaagagatctt |
| EC(up) 1 kbp | atgacgcgaaactggagatccactccccgcgcggtgttcgtttcgtcccgattaatggctttcacaccggg<br>ccgggcaaagtgtctcttgagcatgacgaaatcctcgtcgcctttcattttccgccacagccgaaagaac<br>acgcgggcagcgcgcattttaaatatgccatgcgcgacgcaatggatatttcaacgattggctgcgccgc<br>acattgccgactggataacggcaatttcagcgaattacgcctggcatttggtgttgccgcgccaacgccg<br>attcgctgccaacatgccgaacagactgcacaaaatgcgccattaaacctgcaaacgctggaagctat<br>cagcgaatctgtcctgcaagatgtcgccccgcgttcttcatggcgggccagtaaagagtttcgtctgcatct<br>catccagacgatgaccaaaaaagtgattagcgaagccgtcgccgcggcgggggaaaattgcaatg<br>aatcacagcgaaacaattaccatcgaatgcaccattaacgggatgccttttcagcttcacgccgcacca<br>ggcacgccgctctcggaattactccgcgaacaaggactgctaagtgtcaaacaagggtgctgcgtgggt<br>gaatgtggtgcctgtacggtgttggtcgacggcacagcaatagacagttgcttataccttgccgcctgggc<br>tgaaggaaaagagatccgcacgctggaaggtgaagcgaaaggcggaaaactttctcatgttcagcag<br>gcttatgcgaaatccggcgcagtgcagtgcgggttttgtacgcctggcctgattatggctaccacggcaat<br>gctggcgaaaccacgcgagaagccattaaccattacggaaattcgtcgcggactggcgggaaatcttt<br>gtcgctgcacggggtatcagatgattgtaaatacagttctggattgcgagaaaacgaagtaaaaggatat<br>ccggcctgaattcaggccggattcactg |
| EC(down) 1 kbp | aggttatgtgtttaacaactcatatttcttaatcttgcgatagagcgtagcaatgccgatgcccagttcatcag<br>caacttgcttcttgctgttatgacgtgaaagcgcctcgcggatcatttgcttttccatctcctccagcgccgtgc<br>cgcccgcatcatcgagtgacaggtgcgcctcactgacctctgttacatcactttgctccgttgtgccattattc<br>agcagatttggcggcaatagcgtgctgtcgataacttcacctgaaggaaccacgttaaccagatattcca<br>tcaaattgcttaactcgcgcaggtttccgggccaacgatgcttacgcaatatttcgacgacatcgggagca<br>atgccaggataaaccgatcccagacgacgggtatgcagatgtaaaaagtaatgcaccaatagttcaat |

| | |
|---|---|
| | atcttcctgacgttcacgcagcggtggcagagttatcgggataacattaagtcggtagaagagatcttcgc<br>ggaatttaccttcggcaatgaactgggccaaattctgattagttgcagaaatgatgcgaatgtcgacttgta<br>ttgggctactggcaccaatcggcagaatttcacgtgcctcaatagcgcgcagtaatttagcctgcaacatt<br>aatggcatatcacctatttcatcgagaaacagcgtgcccgtattcgccgcctgaatcaaccctgttttaccg<br>ttggcagaagcgccagtaaatgcacctttaacataaccgaacagttcgctctccagaagctgctccgga<br>atcgcggcacagttgatagcaataaagggtttattccgtcttccgctcaacttatggattgcacgggcgac<br>gacttctttacccgtgccgctttcaccaaccaccataacgctggatgggctgggtgcaatacggctaatga<br>gtcgttttaattgccgcataacacggcactcgccaaccaattgttcaatatgcggttcatcaggtgcattt |
| EC(up) 1.5 kbp | cgtaacgcggtgaagatggctaccggtgttgcaatcaatacactgccgctgacgccaaaacggttatat<br>gaagagttccatctggcaggattgatttgaggataacatcatgtttgattttgcttcttaccatcgcgcagcaa<br>cccttgccgatgccatcaacctgctggctgacaacccgcaggccaaactgctcgccggtggcactgac<br>gtactgattcagctccaccatcacaatgaccgttatcgccatattgttgatattcataatctggcggagctgc<br>ggggaattacgctggcggaagatggctcgctacgtatcggctctgcaacgacatttacccagctaatag<br>aagatcctataactcaacgtcatctcccggcgttatgtgctgcggccacgtccattgctggaccgcagatc<br>cgtaacgtcgctacctacggtggaaatatttgcaacggtgccaccagcgcagattctgccacgccaacg<br>ctaatttatgacgcgaaactggagatccactccccgcgcggtgttcgtttcgtcccgattaatggctttcaca<br>ccgggccgggcaaagtgtctcttgagcatgacgaaatcctcgtcgcctttcattttccgccacagccgaaa<br>gaacacgcgggcagcgcgcattttaaatatgccatgcgcgacgcaatggatatttcaacgattggctgc<br>gccgcacattgccgactggataacggcaatttcagcgaattacgcctggcatttggtgttgccgcgccaa<br>cgccgattcgctgccaacatgccgaacagactgcacaaaatgcgccattaaacctgcaaacgctgga<br>agctatcagcgaatctgtcctgcaagatgtcgccccgcgttcttcatggcgggccagtaaagagtttcgtct<br>gcatctcatccagacgatgaccaaaaaagtgattagcgaagccgtcgccgcggcgggggggaaaattg<br>caatgaatcacagcgaaacaattaccatcgaatgcaccattaacgggatgccttttcagcttcacgccgc<br>accaggcacgccgctctcggaattactccgcgaacaaggactgctaagtgtcaaacaagggtgctgcg<br>tgggtgaatgtggtgcctgtacggtgttggtcgacggcacagcaatagacagttgcttataccttgccgcct<br>gggctgaaggaaaagagatccgcacgctggaaggtgaagcgaaaggcggaaaactttctcatgttca<br>gcaggcttatgcgaaatccggcgcagtgcagtgcgggttttgtacgcctggcctgattatggctaccacgg<br>caatgctggcgaaaccacgcgagaagccattaaccattacggaaattcgtcgcggactggcgggaaa<br>tctttgtcgctgcacggggtatcagatgattgtaaatacagttctggattgcgagaaaacgaagtaaaagg<br>atatccggcctgaattcaggccggattcactg |
| EC(down) 1.5 kbp | aggttatgtgtttaacaactcatatttcttaatcttgcgatagagcgtagcaatgccgatgcccagttcatcag<br>caacttgcttcttgctgttatgacgtgaaagcgcctcgcggatcatttgcttttccatctcctccagcgccgtgc<br>cgcccgcatcatcgagtgacaggtgcgcctcactgacctctgttacatcactttgctccgttgtgccattattc<br>agcagatttggcggcaatagcgtgctgtcgataacttcacctgaaggaaccacgttaaccagatattcca<br>tcaaattgcttaactcgcgcaggtttccgggccaacgatgcttacgcaatatttcgacgacatcgggagca<br>atgccaggataaaccgatcccagacgacgggtatgcagatgtaaaaagtaatgcaccaatagttcaat<br>atcttcctgacgttcacgcagcggtggcagagttatcgggataacattaagtcggtagaagagatcttcgc<br>ggaatttaccttcggcaatgaactgggccaaattctgattagttgcagaaatgatgcgaatgtcgacttgta<br>ttgggctactggcaccaatcggcagaatttcacgtgcctcaatagcgcgcagtaatttagcctgcaacatt<br>aatggcatatcacctatttcatcgagaaacagcgtgcccgtattcgccgcctgaatcaaccctgttttaccg<br>ttggcagaagcgccagtaaatgcacctttaacataaccgaacagttcgctctccagaagctgctccgga<br>atcgcggcacagttgatagcaataaagggtttattccgtcttccgctcaacttatggattgcacgggcgac<br>gacttctttacccgtgccgctttcaccaaccaccataacgctggatgggctgggtgcaatacggctaatga<br>gtcgttttaattgccgcataacacggcactcgccaaccaattgttcaatatgcggttcatcaggtgcatttgct<br>acagaaaaactggtatgcgattggtgaaacgccattaaaaataattgtcggccctgaatgttatgcaattg<br>accaatgattaattcacttttatcgtcccatgaaacaatatgctgcatatgtccatgggtaaaattactctcaa<br>atgttaatggtctgaaacggataggtttcccaataatattattttgcacaacaccaagtgtttttaaggcagtc<br>tgattaacaaactgaacccgattttcatcatctacaactaatacgccctgatccatattatcgatcatggtcg<br>caaatattttactgatgttatctcctggcccctgatcctccagaagtttcgaaacaaaaatggtggatatatg |

| | |
|---|---|
| | gcgaacataatcagaaaattcgcgtaaattatcactgatatgctcttgttgctcgtgggtaacggcaatcaa<br>acttatcaccccaacacaacgatcctgtaaaatgacaggcgtacccagaaatgctttttc |
| EC(up) 2 kbp | ccctgataaaaggccatatcgtgctggttgaacgaccggaagagccgttaatgtcgttaaaagatttggc<br>gatggacgctttctaccaccctgaacgcggcgggcagctctctgctgaaagctccatcaaaccaccac<br>taacccaccggcgtttggctgtacctttgttgatctgacggtcgatattgcgctgtgcaaagtcaccatcaac<br>cgcatcctcaacgttcatgattcagggcatattcttaatccactgctggcagaaggtcaggtacacggcgg<br>aatgggaatgggcattggctgggcgctatttgaagagatgatcatcgatgctaaaagcggcgtggtccgt<br>aaccccaatctgctggattacaaaatgccgaccatgccggatctgccacaactggaaagcgcgttcgtc<br>gaaatcaatgagccgcaatccgcatacggacataagtcactgggtgagccaccaataattcctgttgcc<br>gctgctattcgtaacgcggtgaagatggctaccggtgttgcaatcaatacactgccgctgacgccaaaac<br>ggttatatgaagagttccatctggcaggattgatttgaggataacatcatgtttgattttgcttcttaccatcgc<br>gcagcaacccttgccgatgccatcaacctgctggctgacaacccgcaggccaaactgctcgccggtgg<br>cactgacgtactgattcagctccaccatcacaatgaccgttatcgccatattgttgatattcataatctggcg<br>gagctgcggggaattacgctggcggaagatggctcgctacgtatcggctctgcaacgacatttacccag<br>ctaatagaagatcctataactcaacgtcatctcccggcgttatgtgctgcggccacgtccattgctggacc<br>gcagatccgtaacgtcgctacctacggtggaaatatttgcaacggtgccaccagcgcagattctgccac<br>gccaacgctaatttatgacgcgaaactggagatccactccccgcgcggtgttcgtttcgtcccgattaatg<br>gctttcacaccgggccgggcaaagtgtctcttgagcatgacgaaatcctcgtcgcctttcattttccgccac<br>agccgaaagaacacgcgggcagcgcgcattttaaatatgccatgcgcgacgcaatggatatttcaacg<br>attggctgcgccgcacattgccgactggataacggcaatttcagcgaattacgcctggcatttggtgttgcc<br>gcgccaacgccgattcgctgccaacatgccgaacagactgcacaaaatgcgccattaaacctgcaaa<br>cgctggaagctatcagcgaatctgtcctgcaagatgtcgccccgcgttcttcatggcgggccagtaaaga<br>gtttcgtctgcatctcatccagacgatgaccaaaaaagtgattagcgaagccgtcgccgcggcgggggg<br>aaaattgcaatgaatcacagcgaaacaattaccatcgaatgcaccattaacgggatgccttttcagcttc<br>acgccgcaccaggcacgccgctctcggaattactccgcgaacaaggactgctaagtgtcaaacaagg<br>gtgctgcgtgggtgaatgtggtgcctgtacggtgttggtcgacggcacagcaatagacagttgcttatacct<br>tgccgcctgggctgaaggaaaagagatccgcacgctggaaggtgaagcgaaaggcggaaaactttct<br>catgttcagcaggcttatgcgaaatccggcgcagtgcagtgcgggttttgtacgcctggcctgattatggct<br>accacggcaatgctggcgaaaccacgcgagaagccattaaccattacggaaattcgtcgcggactgg<br>cgggaaatctttgtcgctgcacggggtatcagatgattgtaaatacagttctggattgcgagaaaacgaa<br>gtaaaaggatatccggcctgaattcaggccggattcactg |
| EC(down) 2 kbp | aggttatgtgtttaacaactcatatttcttaatcttgcgatagagcgtagcaatgccgatgcccagttcatcag<br>caacttgcttcttgctgttatgacgtgaaagcgcctcgcggatcatttgcttttccatctcctccagcgccgtgc<br>cgcccgcatcatcgagtgacaggtgcgcctcactgacctctgttacatcactttgctccgttgtgccattattc<br>agcagatttggcggcaatagcgtgctgtcgataacttcacctgaaggaaccacgttaaccagatattcca<br>tcaaattgcttaactcgcgcaggtttccgggccaacgatgcttacgcaatatttcgacgacatcgggagca<br>atgccaggataaaccgatcccagacgacgggtatgcagatgtaaaaagtaatgcaccaatagttcaat<br>atcttcctgacgttcacgcagcggtggcagagttatcgggataacattaagtcggtagaagagatcttcgc<br>ggaatttaccttcggcaatgaactgggccaaattctgattagttgcagaaatgatgcgaatgtcgacttgta<br>ttgggctactggcaccaatcggcagaatttcacgtgcctcaatagcgcgcagtaatttagcctgcaacatt<br>aatggcatatcacctatttcatcgagaaacagcgtgcccgtattcgccgcctgaatcaaccctgttttaccg<br>ttggcagaagcgccagtaaatgcacctttaacataaccgaacagttcgctctccagaagctgctccgga<br>atcgcggcacagttgatagcaataaagggtttattccgtcttccgctcaacttatggattgcacgggcgac<br>gacttctttacccgtgccgctttcaccaaccaccataacgctggatgggctgggtgcaatacggctaatga<br>gtcgttttaattgccgcataacacggcactcgccaaccaattgttcaatatgcggttcatcaggtgcatttgct<br>acagaaaaactggtatgcgattggtgaaacgccattaaaaataattgtcggccctgaatgttatgcaattg<br>accaatgattaattcacttttatcgtcccatgaaacaatatgctgcatatgtccatgggtaaaattactctcaa<br>atgttaatggtctgaaacggataggtttcccaataatattattttgcacaacaccaagtgtttttaaggcagtc<br>tgattaacaaactgaacccgattttcatcatctacaactaatacgccctgatccatattatcgatcatggtcg<br>caaatattttactgatgttatctcctggcccctgatcctccagaagtttcgaaacaaaaatggtggatatatg |

| | |
|---|---|
| | gcgaacataatcagaaaattcgcgtaaattatcactgatatgctcttgttgctcgtgggtaacggcaatcaa acttatcaccccaacacaacgatcctgtaaaatgacaggcgtacccagaaatgctttttcgcggcaatttt ctttactatcgcaaccttcgcaaaggggatcgaagcgagactgtgtcacaactttttcagttttcgtttccagg acgtggcggagcaggcgtgagttgccgctcaactggcgaccaagaaacttcccatacgcgcccgttcc ggcaacgcgacacaagttttcatcaacgatctcaacctcaagctgcaaaacgctggcaagcattctggc aaaacgctgaattgtcggttgaatttgcatcaatactgactgcgtagtagcaagctccatagctttaccttcc agacttacttaaaagtcgatcattgaagacgttgatggttcacagatcatgatgatattaactcaggcgaa attggctttgataaaaacataagatttttatcattttctaatgaaattatggaagagatatcacatttctatatca atatgagaattacggcggtgagtttatcaaactgaagagagatagcctgcccctttat |
| EC(up) 2.5 kbp | aggagatgctaatccgctcacgggcaaacgtatttacagcgcagggttgccggagtgtcttgaaaaagg ccggaaaatctttgaatgggaaaaacgccgtgcagaatgccagaaccagcaaggcaatttgcgccgc ggcgttggcgtcgcctgttttagctacacctctaacacctggcctgtcggcgtagaaatagcaggcgcgc gccttctgatgaatcaggatggaaccatcaacgtgcaaagcggcgcgacggaaatcggtcagggtgc cgacaccgtcttctcgcaaatggtggcagaaaccgtgggggttccggtcagcgacgttcgcgttatttcaa ctcaagataccgacgttacgccgttcgatcccggcgcatttgcctcacgccagagctatgttgccgcgcct gcgctgcgcagtgcggcactattattaaaagagaaaatcatcgctcacgccgcagtcatgctacatcagt cagcgatgaatctgaccctgataaaaggccatatcgtgctggttgaacgaccggaagagccgttaatgt cgttaaaagatttggcgatggacgctttctaccaccctgaacgcggcgggcagctctctgctgaaagctc catcaaaaccaccactaacccaccggcgtttggctgtacctttgttgatctgacggtcgatattgcgctgtg caaagtcaccatcaaccgcatcctcaacgttcatgattcagggcatattcttaatccactgctggcagaag gtcaggtacacggcggaatgggaatgggcattggctgggcgctatttgaagagatgatcatcgatgcta aaagcggcgtggtccgtaaccccaatctgctggattacaaaatgccgaccatgccggatctgccacaa ctggaaagcgcgttcgtcgaaatcaatgagccgcaatccgcatacggacataagtcactgggtgagcc accaataattcctgttgccgctgctattcgtaacgcggtgaagatggctaccggtgttgcaatcaatacact gccgctgacgccaaaacggttatatgaagagttccatctggcaggattgatttgaggataacatcatgtttg attttgcttcttaccatcgcgcagcaacccttgccgatgccatcaacctgctggctgacaacccgcaggcc aaactgctcgccggtggcactgacgtactgattcagctccaccatcacaatgaccgttatcgccatattgtt gatattcataatctggcggagctgcggggaattacgctggcggaagatggctcgctacgtatcggctctg caacgacatttacccagctaatagaagatcctataactcaacgtcatctcccggcgttatgtgctgcggcc acgtccattgctggaccgcagatccgtaacgtcgctacctacggtggaaatatttgcaacggtgccacca gcgcagattctgccacgccaacgctaatttatgacgcgaaactggagatccactccccgcgcggtgttcg tttcgtcccgattaatggctttcacaccgggccgggcaaagtgtctcttgagcatgacgaaatcctcgtcgc ctttcattttccgccacagccgaaagaacacgcgggcagcgcgcattttaaatatgccatgcgcgacgc aatggatatttcaacgattggctgcgccgcacattgccgactggataacggcaatttcagcgaattacgcc tggcatttggtgttgccgcgccaacgccgattcgctgccaacatgccgaacagactgcacaaaatgcgc cattaaacctgcaaacgctggaagctatcagcgaatctgtcctgcaagatgtcgccccgcgttcttcatgg cgggccagtaaagagtttcgtctgcatctcatccagacgatgaccaaaaaagtgattagcgaagccgtc gccgcggcgggggaaaattgcaatgaatcacagcgaaacaattaccatcgaatgcaccattaacgg gatgccttttcagcttcacgccgcaccaggcacgccgctctcggaattactccgcgaacaaggactgcta agtgtcaaacaagggtgctgcgtgggtgaatgtggtgcctgtacggtgttggtcgacggcacagcaata gacagttgcttataccttgccgcctgggctgaaggaaaagagatccgcacgctggaaggtgaagcgaa aggcggaaaactttctcatgttcagcaggcttatgcgaaatccggcgcagtgcagtgcgggttttgtacgc ctggcctgattatggctaccacggcaatgctggcgaaaccacgcgagaagccattaaccattacggaa attcgtcgcggactggcgggaaatctttgtcgctgcacggggtatcagatgattgtaaatacagttctggatt gcgagaaaacgaagtaaaaggatatccggcctgaattcaggccggattcactg |
| EC(down) 2.5 kbp | aggttatgtgtttaacaactcatatttcttaatcttgcgatagagcgtagcaatgccgatgcccagttcatcag caacttgcttcttgctgttatgacgtgaaagcgcctcgcggatcatttgcttttccatctcctccagcgccgtgc cgcccgcatcatcgagtgacaggtgcgcctcactgacctctgttacatcactttgctccgttgtgccattattc agcagatttggcggcaatagcgtgctgtcgataacttcacctgaaggaaccacgttaaccagatattcca tcaaattgcttaactcgcgcaggtttccgggccaacgatgcttacgcaatatttcgacgacatcgggagca |

| | |
|---|---|
| | atgccaggataaaccgatcccagacgacgggtatgcagatgtaaaaagtaatgcaccaatagttcaat<br>atcttcctgacgttcacgcagcggtggcagagttatcgggataacattaagtcggtagaagagatcttcgc<br>ggaatttaccttcggcaatgaactgggccaaattctgattagttgcagaaatgatgcgaatgtcgacttgta<br>ttgggctactggcaccaatcggcagaatttcacgtgcctcaatagcgcgcagtaatttagcctgcaacatt<br>aatggcatatcacctatttcatcgagaaacagcgtgcccgtattcgccgcctgaatcaaccctgttttaccg<br>ttggcagaagcgccagtaaatgcacctttaacataaccgaacagttcgctctccagaagctgctccgga<br>atcgcggcacagttgatagcaataaagggtttattccgtcttccgctcaacttatggattgcacgggcgac<br>gacttctttacccgtgccgctttcaccaaccaccataacgctggatgggctgggtgcaatacggctaatga<br>gtcgttttaattgccgcataacacggcactcgccaaccaattgttcaatatgcggttcatcaggtgcatttgct<br>acagaaaaactggtatgcgattggtgaaacgccattaaaaataattgtcggccctgaatgttatgcaattg<br>accaatgattaattcacttttatcgtcccatgaaacaatatgctgcatatgtccatgggtaaaattactctcaa<br>atgttaatggtctgaaacggataggtttcccaataatattattttgcacaacaccaagtgtttttaaggcagtc<br>tgattaacaaactgaacccgattttcatcatctacaactaatacgccctgatccatattatcgatcatggtcg<br>caaatattttactgatgttatctcctggcccctgatcctccagaagtttcgaaacaaaaatggtggatatatg<br>gcgaacataatcagaaaattcgcgtaaattatcactgatatgctcttgttgctcgtgggtaacggcaatcaa<br>acttatcaccccaacacaacgatcctgtaaaatgacaggcgtacccagaaatgctttttcgcggcaatttt<br>ctttactatcgcaaccttcgcaaaggggatcgaagcgagactgtgtcacaactttttcagttttcgtttccagg<br>acgtggcggagcaggcgtgagttgccgctcaactggcgaccaagaaacttcccatacgcgcccgttcc<br>ggcaacgcgacacaagttttcatcaacgatctcaacctcaagctgcaaaacgctggcaagcattctggc<br>aaaacgctgaattgtcggttgaatttgcatcaatactgactgcgtagtagcaagctccatagctttaccttcc<br>agacttacttaaaagtcgatcattgaagacgttgatggttcacagatcatgatgatattaactcaggcgaa<br>attggctttgataaaaacataagatttttatcattttctaatgaaattatggaagagatatcacatttctatatca<br>atatgagaattacggcggtgagtttatcaaactgaagagagatagcctgcccctttatcttatttctgatactt<br>agcagcaaataaataacgcgataaaaaaagccaaacgttttcgtattttacaaacaaccagaagctgg<br>catcaatttgtgatcaaccccacacattatccgtcaaattagtcttttgcagccgcgcggataattctggcac<br>acttattgttagtcccaggtatagctgtgaaaacaccaatcactttggcaagtcacagtgaaataaaccac<br>tttgcctgtcattccactaccgggactttatgatgaaaactgttaatgagctgattaaggatatcaattcgctg<br>acctctcaccttcacgagaaagatttttttgttaacgtgggaacagacgccagatgaactgaaacaagtac<br>tggacgttgccgcagcattaaaagcactgcgtgctgaaaacatctcaaccaaagtctttaatagtggatta<br>ggtatttccgtattccgcgacaactccacccgtacccgcttctcttatgcttccgcg |
| ST(up) 2.5 kbp | acgtagcagcaggggtatcaacgtttgcatttcaaggtgccgggcttcccgtcctacgctggtaccctgct<br>cttgcgttaatttttggtggcacatatcaagcgcctcaacagccttcgccgccgctttgtcaacaaggtgcgt<br>aagattgctgcgggttaacggatctaacgtacagccaaagttatgttcaatgcagctggcaatatagggc<br>atcacctcctgcataacaagattcgtcgataatttacttaattcaccgccagtgttatttttgataatatctaac<br>agctgcttttccaggttttccagcttcgcttccgctttctttgtttctggcagccatggcccaaaagctgacttttct<br>ttcaggccatcttttatgatttgcgcggtatactctgcccccaccttcatcagtagcgtcttcgcctcaggaga<br>atcactggtggcgttgagcgctgaacgaaagagcccggcaaactccattatcgctttcttaccggcgaca<br>ttatttgaattggtaaaaacttcttttaacgcctcagcgtctttcccgcatttaaacaatgcatccagactcgcc<br>tgtttgatcagcgcgggaaaatcttccagttgcgggcctttaatttcccctgacagcgtcgctgtggcactttc<br>tctgactgcggaaagattcgccgcaagattcgtggcctgcgttttgatctcggtctgcatacctggcattatg<br>acggggggctgagtccttacacttgtaaccattattaatatcctcttctgttatccttgcaggaagcttttggcg<br>gtttccaggctgctacttatcgtactgctcagcacttttaccaggttgtcgtacaatgaattggcattgctatatt<br>tttgcgtcagcgtctgtaatgtggttttcatattttcttcctgcgctttaaaacccgactgccaggcttgatatttg<br>gcgttatccatttcgagttttgagtcttttcccggcgcgcctaaaccatcaatatcctgaaccatttttttgtaatg<br>gcgtcagatcaacggtgacgacataaccggatccataagatttcaggcagctattcggtaaattcaattc<br>actgagccactgtctcgcttccgcttcagtggctactttaacgccgctgcctgactgcgctggaaataaaac<br>ggtattactgtttatttgattatatttattgactaaactgtttaaatcatttttgagtgaggtaacatctagcttaacg<br>gtattaccgtccttacctggtaataaccagcctcccattttggaaagaatatcactgaaggcctgataaaa<br>atcggtatagactgcgacaacgttttcataaacgcccagatagctgtcacctatcgccgatatattttggga<br>aaccatatcccaaatctcagcatcagaaatggttgttctcggctgcgccataggcgaagcgctaaataag |

| | |
|---|---|
| | gccgacgtcggcgcagaaaacgcgctccgcaggttctcattttgttctgcggataatgacacgccggact<br>tcgccagcgcattcaggctgctggtcaactgctggcgcgccagcgtgcgctcgtcattattctcttcagaga<br>tcggtggcgttgactgcagcgtctgctgtgcctggtggattttagtagccgcctgcgataatgaaatgatatc<br>tgtaccgcgatgttctgtggtagacggtaccacggcagtctcgacgtgctcgctcgccgagggagtctgc<br>ggccgttcggcaacgatccccggatgaggagaagcggaataattttgaatattaagcataatatcccca<br>gttcgccatcaggagcgcgattaaatcacacccatgatggcgtatagatgacctttcagattaagcgcga<br>atattgcctgcgatagcagcgagtgcggatgctttcgactggttaatgctctccattgttttcagcatttcctga<br>atcaggctggtcgatttacgtgaactttcacgggcttcgtccgatgcggtgctggcaacccggttattcacct<br>ggctaatttgctgctcggaacgttcctgagtagcggcgtactgcccggacgcccctgcaataccaccgac<br>cgtgaccgagttcttcataatcagatcgcccgtcatctgcatcttgcgcgcatcgattcgggtcatatccatg<br>gtattctgctcaagacgaatatcggattcgacagactcaagacgtttcgacagaatagcctgatgttcagg<br>ggagatttgtttattactgtctttaatacccagactttccgtggcgctggttccggcattagatttaagcgtcgc<br>atcattaagatttttcgtcgcatcggtaccggttttcttcatatttaacgatttcagagaatcgacgccttcagc<br>accgagtttgacgctattctgcccgttcagcacgtttttaatactgtggctttcagtggtcagtttatcgatcttcg<br>cggcattatgtttaag |
| ST(down) 2.5 kbp | cgcgcctctttcattctgcagccccttatattccagtttggcgcccacgccagtgatccccaactgaagcgc<br>gctctgggaaatactaccggacaacgcattcatcccttcgcgcatcatggagcttgccgtcgttttagctgc<br>atcaaaactgactaatgacaacttaccagacagtttgctatcagcctggttcaacgtcagcattaacgtatt<br>cgcggcagccaacagcgcaacggcactggaagacattccgctaatatcaaaaaactttccgacttctg<br>cctgctgctcgcgtaactgggtttgcacaacctcattcgctttagtcgtgacattatttgccagagcattcaaa<br>tcctgattcatgtcggtattttgaatactggcttttaaaaaggacgtgatcgttccgggggtttgcgttaatacc<br>cctggcgcaggcgcgctcagtgtaggactcaaccccaggtcactgactttactgctgctaataccaatact<br>attcagaatatctttagcgctaacggattgcgaagctgtctgtgaactattctcaacagaatgattatttaaat<br>aagcggcgggatttattcccacattactaattaacatatttttctccctttattttggcagtttttatgcgcgactct<br>ggcgcagaataaaacgcgaagcatccgcattttgctgtaccgcagaagacatggctttttgcagttccgc<br>cgttaccttctggttttcaccaaatatttctacggattgtttaagccactgctgaatctgatccatggcaaaacg<br>ggcgagcataaaatcagcaagcgcctcgctggcatttttaataaatacgccctcggcaacaccaccgg<br>ctgactgggctgcggtattcgtgacttccatgcccaacgccactttatttagggtattacctaccagctcttta<br>cttaaggcattcgtttgcaggcccatcttgctacccacattacccagaccgctagtaatacgttgcatcccct<br>gggtaaagagtttgctgccgttttgcgccaactgtttcagcacgttaggcaccaacttcttaatcgtttcgccc<br>atcattttgctcagcgcgttacccagtttcgccgccgcgcctttcccgacaactgcgaccaccacaatgac<br>cgccaccatggcaatagcggcgacaatcgcaccaacaatgctgccggccatctctgccgttttcttatcg<br>acgcctaatccttccagcgctttggtaatcgccttgccaatcagctccattaacggcttcagcacatgctcc<br>ataatcgggtttagcgcctgctgaataaacgacactcccgtcgccgccttcacaatttcatcggccaccatt<br>accgcaagtcccaccgcagccagcgccagactcgccccaccggtaaaaacagcggccacaacgct<br>gacaatggttagcagcgcgccgaggactttcccgatacatcccataatgcggttcgtttcctcggctttgcg<br>cgtctcttcctggaattcagccgatttcttttccatctccgcctgacgcccttcctgcaaggcgttgaaaagcg<br>caagatcgttttgcaggctttcttccgtatttttgcccacaatctcaataaacatggccatgagcatagtgag<br>gcgggcgacatttgacagattatcctgctcaccctgggaaacctgattctgagaggcggcattagccgttc<br>cctggaatttggtcagaatgttatccgctttctcggctttcgctttggcgtctgtgcctgctttaaccgtcgcatc<br>cgtggccttatctaaggcctctttcgcctctgtcgcttcttttccggcctgttctaccgcggcttcagcttgtgcat<br>agccggggtcagccgggtccagcgattgcaatttattttgcgcctgcgtcagtttttttggtcgcagcgtcata<br>aacactcttggcggtatccgtctttttgatactggcttcatagagatccgtcgcctcctgagcctctcccagag<br>ccgtctggaattctttcgatacctgaatccccatctctttttgtgactcaatcatcgcctgccataccgccaga<br>cgagactccagttgagacagcgaaacatcgcccagtagggtcattaacttgccaagcagtaatgtcaat<br>tgcccttcgctggagagtttttcccgggcggcgtccgtaggcggctttagacccaccgtattaatagcgctc<br>tcgccggactttgttccggctttaaggtcgcccgctttcgttgccaccacatctttaaaagctttatccgccgct<br>tttaaaaagtccgtgttcttacgaacgccttcaaaagccgcctcagcgaggcgcggattttgggtatatccg<br>ctacggctaatgctacttgcgtcatttaccataattattccttttcttgttcactgtgctgctctgtctccgccgttttt<br>agcgcctccagatagaccaacgcttttg |

| | |
|---|---|
| SA(up) 2.5 kbp | ctatcgtgtcatgtaatcttgcatccgatcttgcaacgctgtaaatgtttcgaagccatcctcttctaagaagtg<br>ccctccatcttccacgattcgcaagttcccctctaatgcattcattaaacgctgggtttctttatatgaaacgta<br>tttgtcatttttagaactcaatccgtaaaaattgtcaactttctttttaatattatcgtaatcaatggttacattactt<br>aaatcaatatctaaatctatattttctgcatcttctttaaagcccgctatactaaaaaagccttcaatcggctg<br>atcaatcatttcaatatattttaaagctgtgattgaacctaaaccatgtgttacaaaatatgtatcctttttgcgt<br>acattaatttgtttcgtcatagcttcaatccactgatccactgtcttcgcttcaggggattcaaaattaaataat<br>gttacgtcatatccttctaaagttaagttatgctccaaccactgataccaatgatttctactatttccatgcatag<br>aatgtacaataattacatctgtcatctcattctctcctttcaacttactacttcttttctatttttaaaaaaatgactg<br>attacctataattgtaaaataaaaacaccttaattagaaatgttatatcgcaaagtgacatttctaattaaagt<br>gtattgtcatcatttcaatatcattcaaaaacagctaaacctttgtctctgcttcaatttcacaaaaataattccc<br>gctgaaagtatctatatttacacattacttccaccattatataacttaaaaatgactatatttcatcaaacattat<br>ctaaaggcgtcgcacctacaccaacaccatccaacaattaacttacaactctgcgattacttcttcagcag<br>caactttaccttgtgtaatacaatcaggtagtccaaccgcttcaaaagatgcaccagttactctaagtcgtg<br>gatatgtttgtttaatatgtgcttgaatctgtctaatttgttgaatatgaccgacatggtactgtggcatacttttcg<br>gcaaacgattgacaattgtaaattcaggatcacctttaaatgtcatcatttgacttaaatctctacgtacaatc<br>gatactaattcattatctgtatgatcatcaaccacagtatcacctggtttacctacatacgcacgaatcaaaa<br>ccttaccttctggtgtagtaaatggccatttttcgatgtccaagtacatgcggtaatgtctgtatcactcgttctc<br>gcaatcacgaagccagtaccatcataagtattttcaatgtctttttcatcaaatgccaatacaacagttgca<br>acagtcgtactatccatcgttttaaagtaatcaaatgctggatcttgtccgaaccaattcaaaaagacttgat<br>gcggtgttgtcactaataccccatcgaatacatcttcttgttgattactgtaaacaattttatattgcttttgagat<br>gtaataatatcatccactgacgtattgtagcgtattgtcacacctttatttttcacatcttgttctaatgcttcaata<br>aatgagcttaaaccatgcttaaattgtttgaattgtccttttggtgcgccaggatataattgtctttgtttcagac<br>gcttatttttctcatccttcataccttttatcagacttccgaatgcctcttctttttctttaaaattaggaaacgtactc<br>atcaaacttaatttatcaatatcggtaccataaataccacccattaaaggctcaattaagttctcaagtacct<br>cattacctaatcttgctctgaaaaatgcaccaacagaaatgtcaccatcttgcatttgtataggctttttgatta<br>aatctaatcctgctcttaatttaccaagtggcgatattaattttgtagtaacaaatggtttaatatctgttggaat<br>acccataattgaaccacctggaatcggatataatttatttttcgcaaaaatatatgattgtccagtcgtatttgt<br>aacaatatcttgttctaatccaatatctttcgctaattctgtcataatcgtttttctacctaaataagattcaggcc<br>ctagttcaatcatataaccatctttacgatacgattgaatctttcccccggacgattcgatgcttcaaagatg<br>gttacatcaatattaggatcttgctgttttaa |
| SA(down) 2.5 kbp | gacttgatttcatcaacaattgcaccgataaataatggatgtgtattcggcatttttggacgataataattcgc<br>accaatatcatcgcaaacaactttacattcataatcattgtcataaagcacctctaaatgctcacatacaaa<br>acctactggcgtatatataaagtttttatactgatgtttttcatataaatcacgtgttaaatcttgtacatctggcc<br>ctaaccaaggtgtacctgtattaccttcagattgccaaccaatcgcgatatgttcaatattagattgttctttaa<br>ttaaaagcgcagtatgttctagttcttgtggatatggatcattattcttttcgattaaacctttggcaaactatgt<br>gccgaaacaactaataccgtgtctttatgttcctcttccggtatttgagctaatgtttcgttgactttattcgtcca<br>atattcaataaatttaggttgttcataataatgtttcacatgtgtaagttgaataccatattttgcagcttcttcatc<br>agcacgtttgtcatatgatcctactgaaaatgaagaataatgtggtgctagtactaccgtgattgcttcagta<br>ataccatcattgtgcatttgttcaaccgcatcttcgataaatggtgaaatgtgttttaatcctaagtatagtttaa<br>attcaacatctgcatatgctttatttaatgctgaaactagtgcatcagcttggtcatctgttgtacctgctaatgg<br>tgataaaccacctataaattcatatctatctttcaaatcttgaagttcttcttcagatggacgtttaccatgtcta<br>atatctgtataatatggctctatgtcactttctttataaggtgtgccataagccataactaataaccccattttttt<br>agtcattgataataccttcctttaaatgaattatctttcatgtgcttcaatgtaatactatgattatctttgtgtatat<br>gtgtgtacgaattcgcttactttacgtaacgtctctggttgcacttctgggaaaacaccgtgtcctaaattaaa<br>gatgtgtttaccgttctccataccttgatctaatattggtttcaatctctcttcaatgacattccatggtgctaata<br>aaattgatggatctaaattcccttgtaatgttttagtaacgcctaattgttgagcctgattaatagacgttctcca<br>atctaggcctaatacatcaatcggtaaatcattccattcattgattaaatgactggcacctacaccgaataa<br>aattaccggcacatcatgtttttctttaacctcactgattaatcgaatcatatgtggtttaatgtaacgtctgtaat<br>cctcgacatttaatgcacctacccatgaatcgaaaatttgaatcaattcggcacctgcttcgacttgagctgt<br>tacatatttaacagatacatcaactaaatgattcattaaagcaaaccatgttgcttcatctctatacatcatcg |

| | |
|---|---|
| | cttttgtaaaattgtaatttttcgatggtccgccttcaatcatatatgacgctaatgtaaatggtgccccagtaa<br>atcctattagcggcacatttaacttttcttctgttaaaagtttaattgtatctaatacatatggtacatctcgttcgg<br>ggtctatttgagaaagtttctcaacatcttgaattgttttgataggattatgaatcactggaccaatacccgatt<br>taatttctacatcgacaccaattggctttaatggtgtcataatatctttgtataaaattgctgcatctgtatgata<br>attatcaactggtaaatgtgttacataagcgcacaactccggctgatgtgtaatatcgaatagtgaatattttt<br>ctttcaattttcgatattctggttgcgaacggccagcttgtcgcataaaccaaacaggtgtatgtgatgtttctt<br>cacctttgatcatttttaaaattgtattgtttttattatgcaccataaaggcctcctaaattaaaatcattcttatcta<br>tattatcatatcgctcattcgttcgtattttcaataaataaatgtcataaaactgacatttaatcatagaactattt<br>attgtaaatttaaattctaaagtccattattttgtatcattacttctaaatatctcgcaagattcattatagtaatttt<br>aatcaattattaatagtggtaatgactagtttatcatcgtataataaataaaaacataaggggggacctttcat<br>atgaagaaactatatacatcttatggcacttatggatttttacatcaaataaaaatcaataacccgacccat<br>caactattccaattttcagcatcagatacttcagttatttttgaagaaactgatggtgagactgttttaaaatca<br>ccttcaatatatgaagttattaaagaaattggtgaattcagtgaacatcatttctattgtgcaatcttcattccttc<br>aacagaagatcatgcatatcaacttgaaaagaaactgattagtgtagacgataatttcagaaactttggtg<br>gctttaaaagctatcgtttgttaagacctgctaaaggtacaacatataaaatttatttcggatttgctgatcga<br>catgcatacgaagactttaagcaatctgatgcctttaatgaccatttttcaaaagacgcattaagtcattactt<br>tggttcaagcggacaacattcaagttattttgaaagatatctatacccaataaaagaatag |
| CD(up) 2.5 kbp | atgaagcaatatatagtcattgggtgtgggagatttggaagttcagttgcgtctactatgcatcttttaggaca<br>tcaagtaatggcaatagacaaaaatgaagattcagttcaaagtatatctgacaaggtaacccattcactt<br>atagtggatgttactgatgagcaagcgttaaggtcattaggtttaggtaactttgatgtagcagtagttgcaa<br>taggttctgatataagggcatctataatggcgactcttatagccaaagaaatgggtgtagagttgataatat<br>gtaaggcaaaggatgaattacaagctaaagtgctttataaaattggcgcagatagagttgtatttccagaa<br>agagatatgggagtaagagttgcacacaatttagtttcggataatatattagaccatattgaacttgaccca<br>gagtattcaattgttgaaatcgtaactccaaatagttgggttggcaagacacttatagagcttgaattaaga<br>gctagatatgagataactgtacttgctataaaaacaggtaaaaatataaatgttacaccttctccagatga<br>ggaacttacagccggaagtatcctagttataatcggtcaaaatactagtataacagcgataacatctgga<br>aataaggggataattagaagaagataatttactatttaatatatatttaattgcaatgaaagtaaagagtatc<br>atataattatgaatagttatatgatactatttttatttaatcgaaggtagtatttatttgtaagattagataaagag<br>aagttaaattaaaagtaaggaggctgtgctaatataaaaatttataagttattagcatttagataaaatgatt<br>acaaatataaacagtaaggataatgaaaagttaaagtatacaagagcactattaaaatcaaagaatag<br>gaataaagagtcaaagttcataatagaaggatacagaatagtaatgcttgcacttgaatgtatggcaaac<br>cttgattatgtatttatcaatgaagaatttgaaaataagaaagaacatgtaaaactattagaagatttggata<br>aaaaaaacacaaagatatacaagactactaataaaaactttaaagaattagtggatacagaaaatact<br>caaggaataataggtgtagtttcatttaagaaaaaaaaattaagtgaaagtataaataaaaaagataaa<br>tttgtattgattttagatagaatacaagacccaggaaatatggggactataataaggactgctgattctgctg<br>gagtagatgccataatagcactaaagggatgtgtcgatatatacaacccaaaagtaattaggtctactat<br>gggttctatttttgatatgaatataattgatgcttcacaagatgaaactgtggacatgcttaaatcattggatttt<br>aatatagtttcaagttacttaaatacagaaaatttttatgacaaaatagattatggttcaaaagtagcattggt<br>gataggaaacgaagcaaatggaataaatgaagaacttgtatcaaagtctgatattttggttaagatacct<br>atatatggtaaagccgagtcgttaaatgctgcgataagctctgctatactgatgtatgaaataaaaaaata<br>cttaatttaatgtattgtaataaatataatgtcatgttataatcttgaattaaatagtagagtatcaaaaatagtc<br>aatatatattaaaaaaataattaaattaattatatattagatgtatatcatataaaaaatgattgagttgattat<br>gtattagatatatattatataaaaaataattgagttaattatgcattagatgtatattatataaaaaataattga<br>gttaattatgcattagatatatattatataaaaaataattaaattaattatgtattggatatatattaaaaatagt<br>cacataatttgaatggaaatgatataatactaaaataaacaatataatattgtaaatgcaatgaaagagg<br>aaagtattttgattaaacttagtaaagagataaacacctaggctgggagtgttttctaagaggtcatgaga<br>agttccctctggagtaacagagctgaaattttacagtaggctttgacgtcaaaaacgcgttaagtttgttag<br>aggtggtttgatgatttttaaattgttaaactactagggtggtaccgcgaaactata |
| CD(down) 2.5 kbp | tagacagggattgaggggctttttttatacaaaaaaacgaaagggtgatatgtgtgcaagaaaaattactt<br>gctttacgtgaagcagctttggctgaaataaaagaagcacaaagcatagaaagtgtagaaagtttaaga |

| | |
|---|---|
| | gttaagtacttaggaaaaaaaggtgagataactgccatacttaaagaaatgggtaaattatctgctgaag<br>aaagaccagtagttggtaaggttgccaatgaggtaagagaaaacattgaacttagcataaattctaaaa<br>aagaagaaataaatgctattgaaaaagaaagaaaattaaaagaggaagtgatagatgttactcaacc<br>aggaaaagttttaaaggttgggaagaagcatccaataactcaaattatagatgaagtaacagatatattt<br>atcggaatgggattctctatagcagaagggccagaagttgagactgttgaaaacaactttgacgcattaa<br>acgctcctaaagaccatccatcaagagatatgagcgatacattctatatcaatgatggggtattacttaga<br>actcaaacatctccagttcaagtaagaactatgagaagtcaagagttaccaataaaagtaattgcacca<br>ggtagatgttttaggtcagactcgccagatgctacacactcaccaatgttccatcagatagaagggcttgtt<br>gttggaaaagatgttactatggcagaatttaaaggaactatggatatcttcgttgaaaaattgtttggttctga<br>tatcaaaactaagtttagacctcacaacttcccatttacagaaccaagtgcagaggttgatgttacttgtttc<br>aaatgtggtggtaaaggttgcccaatgtgtaaatatgaaggttggatagaaatattaggttcaggtatggtt<br>catccaaatgtgcttagaaattgtggaatagacccagaagtttacagtggatttgcatttggagttggggttg<br>aaagacttgcaatgcttaaatacgaaatagatgatattagattattattcgaaaatgatatgagattcttaaa<br>tcaattttaattaggagggtatgtagatgttagtatctttaaaatggcttagagactatgttgatatagacatgg<br>atgtaaaagagttcgctgataaaatgacaatgacaggaactaaagttgaaacaatagattattatggtga<br>agaaatagaaaatatattggttggaaagattttagaaataaaacaacatccaaatgctgataagttggttg<br>taactaaagtagatattggagataaagttgttcaaatagttacaggagctacaaatatatcagaaggaga<br>ttatattccagtagctgtaaatggttctaagttacctggaggagttgaaatcaaacagactgatttcagaggt<br>gaattatcagatggtatgatgtgttcagcagctgaactaggtatagatgaacattacattgaggagtataa<br>aagaggtggtatatatattttagaccacgaagattcttatgaattaggaaaagatataaaagatgttttagg<br>attaaaagatgctttaatagattttgaattaacttcaaacagacctgattgtaaatgcatgatgggtatagcta<br>gagaagcagctgcaactataggaacaaaagtaaaatatcctgaaatcgaagtaaaagaaagtgacg<br>aagagatagatttcaaagttgagatagataatccagatttatgtagaagatatgttgctagaatggttacag<br>atgtaaaaatagaaccttctccatattggatgcaaagaagacttacagaagcaggagtaagacctataa<br>gtaacatagtcgatataacaaacttcgtaatgttagagcttggtcaaccacttcatgcttttgatataaatcaa<br>gtagagactggaagaatagtagtaagaaatgctaaagatggagagaaacttgtaacattagatgatgtt<br>gagagaacattagataaagatatgctagttataacaaatggagaaaaatcacttggtttagctggtgtaat<br>gggtggtgctaactcagaaataacttctaatacgaagactgtactttttgaaagtgccaatttcaaaccaga<br>aaacataagaatgacagctaaaaaagttggtattaggtcagaagcatcttcaagaaatgaaaaagactt<br>agaccctaatcttgcagagatagcagcaaatagagctgcacaacttgttgaaatgttaggagcaggaa<br>aagttttaaaaggtgttgtagatgtatatccaaataaaccagaacctaaaaaattggtagtaaatcctcaa<br>agaattaaccacctattaggtgtagatgtaccaatggagcagtttgtaggaattttagaatcattagagttta<br>aatgtaatttggtagctaatgataaattagaaatagatgtaccaagctttagaacagatatggaacaaga<br>agctgatgtatgggaagaaatagctagaatttatggatttgagaata |
